# Cdc42 GTPase Activating Proteins (GAPs) Maintain Generational Inheritance of Cell Polarity and Cell Shape in Fission Yeast

**DOI:** 10.1101/2020.06.16.151308

**Authors:** Marbelys Rodriguez Pino, Illyce Nuñez, Chuan Chen, Maitreyi E. Das, David J. Wiley, Gennaro D’Urso, Peter Buchwald, Dimitrios Vavylonis, Fulvia Verde

## Abstract

The highly conserved small GTPase Cdc42 regulates polarized cell growth and morphogenesis from yeast to humans. We previously reported that Cdc42 activation exhibits oscillatory dynamics in *Schizosaccharomyces pombe* cells. Mathematical modeling suggests that this dynamic behavior enables a variety of symmetric and asymmetric Cdc42 distributions to coexist in cell populations. For individual wild type cells, however, growth follows a stereotypical pattern where Cdc42 distribution is initially asymmetrical in young daughter cells and becomes more symmetrical as cell volume increases, enabling bipolar growth activation. To explore whether different states of Cdc42 activation are possible in a biological context, we examined *S. pombe rga4*Δ mutant cells, lacking the Cdc42 GTPase activating protein (GAP) Rga4. We found that monopolar *rga4*Δ mother cells divide asymmetrically leading to the emergence of both symmetric and asymmetric Cdc42 distributions in *rga4*Δ daughter cells. Using genetic screening approaches to identify mutants that alter the *rga4*Δ phenotype, we tested the predictions of different computational models that reproduce the unequal fate of daughter cells. We found experimentally that the unequal distribution of active Cdc42 GTPase in daughter cells is consistent with an unequal inheritance of another Cdc42 GAP, Rga6, in the two daughter cells. Our findings highlight the crucial role of Cdc42 GAP protein localization in determining the morphological fate of cell progeny and ensuring consistent Cdc42 activation and growth patterns across generations.

## Introduction

Symmetry breaking is an important process that enables cell polarization and promotes essential cellular functions such as morphogenesis, cell migration, and asymmetric cell division (Johnson et al., 2011; Turing, 1952). Disruption of these functions impairs normal development and cellular differentiation, fostering the onset of disease (Geiger and Zheng, 2014; MacDonald, 2014).

The conserved GTPase Cdc42 is a master regulator of polarized cell growth in eukaryotes ranging from yeast to humans (Etienne-Manneville, 2004; Johnson, 1999). Targets of active Cdc42 promote growth by enabling functions such as cytoskeletal organization, membrane recycling, and polarized secretion (Etienne-Manneville, 2004; Heasman and Ridley, 2008; Perez and Rincon, 2010). Transitions in Cdc42 activity are regulated by guanine-nucleotide exchange factors (GEFs) that promote the activation of Cdc42, GTPase activating proteins (GAPs) that inhibit Cdc42 by accelerating GTP hydrolysis, and guanine dissociation inhibitors (GDIs) that extract GDP-Cdc42 from the cell membrane (Cole et al., 2007; Koch et al., 1997; Sinha and Yang, 2008). Once activated, Cdc42 associates with cell membranes via its prenylated C-terminus (Choy et al., 1999), where it then undergoes autocatalytic activation thus enabling symmetry breaking (Butty et al., 2002; Irazoqui et al., 2003; Kozubowski et al., 2008).

Several experimental as well as mathematical modeling studies have examined the mechanism of Cdc42 amplification and symmetry breaking during bud-site selection in the budding yeast *S. cerevisiae* (Chiou et al., 2018; Irazoqui et al., 2003; Kozubowski et al., 2008; Slaughter et al., 2009; Wedlich-Soldner et al., 2004). In the model of Goryachev et al., this is realized through a winner-takes-all mechanism that ensures selection of a single bud site (Goryachev and Pokhilko, 2008). These studies suggest that stochastic activation of Cdc42 at the cell membrane recruits a GEF-associated complex consisting of the scaffold protein Bem1 and the GEF Cdc24, which then promotes further activation of Cdc42 in a positive feedback loop (Bender and Pringle, 1989; Chant and Herskowitz, 1991). The emergence of activated Cdc42 clusters exhibits characteristics of activator-inhibitor Turing models. These models rely on slow diffusion of an activator coupled with fast diffusion of an inhibitor (Bendezu et al., 2015; Goryachev and Pokhilko, 2008), thus enabling concentration of the activator and inhibition of nearby activator clusters. In this self-organizing system, symmetry breaking can occur in the absence of polarity landmarks as a dynamic process that initially consists of multiple small clusters of active Cdc42 that compete with one another for regulators until the largest cluster with the most robust positive feedback eventually wins (Howell et al., 2009).

In many complex cell types, such as neurons, symmetry breaking is followed by the organization of multiple sites of growth. An excellent model system for the study of symmetry breaking that allows polarization of multiple sites of Cdc42 activation is the fission yeast *Schizosaccharomyces pombe*. These cells are rod-shaped with two defined growth zones at the cell poles, which allows for straightforward measurement of changes in cell polarity, growth, and cellular dimensions. In addition, the life cycle of *S. pombe* cells includes both symmetric and asymmetric states of growth. *S. pombe* cells divide in the cell center with each new daughter cell inheriting one cell tip that was previously growing (Mitchison and Nurse, 1985; Streiblova and Wolf, 1972). Upon cell division, the old cell end (OE) activates Cdc42-dependent growth first in a process termed old-end takeoff (OETO) (Mitchison and Nurse, 1985; Streiblova and Wolf, 1972). The cell continues in this asymmetric growth pattern until it reaches a length of approximately 9 μM, when growth begins in the new end (NE) of the cell in a process called new-end takeoff (NETO) (Mitchison and Nurse, 1985). During this process, the activation of Cdc42 follows a similar pattern and can be visualized using the CRIB-GFP bioreporter, which consists of a GFP-tagged CRIB domain that binds active GTP-Cdc42 (Tatebe et al., 2008). CRIB-GFP is initially present at the NE as a remnant from the cell division site, but it quickly disappears from the NE and a patch of GTP-Cdc42 emerges in the growing OE (Das et al., 2012). Although small amounts of CRIB-GFP can be observed in the NE before NETO, growth is not detectable until the fraction of CRIB-GFP found on the NE exceeds approximately 20% of the total CRIB-GFP found on the cell cortex (Das et al., 2012). After that threshold is reached, NETO takes place and cell growth begins from the NE (Das et al., 2012).

We previously showed that the activation of Cdc42 in *S. pombe* exhibits oscillations that are anticorrelated between the two cell tips (Das et al., 2012). Mathematical modeling described this oscillatory behavior as resulting from both positive and delayed negative feedbacks, as well as competition for active Cdc42 or its regulators (Das et al., 2012). One of these Cdc42 targets, the Cdc42 GEF Scd1 (bound to the scaffold protein Scd2), generates a positive feedback loop of Cdc42 activation, analogous to the Cdc24–Bem1 complex in *S. cerevisiae* (Chang et al., 1994; Das et al., 2012; Endo et al., 2003; Wheatley and Rittinger, 2005). Another target of Cdc42 in this complex, the PAK kinase Pak1 (also known as Orb2 and Shk1), functions as a negative regulator of Cdc42, generating a negative feedback loop (Chang et al., 1999; Das et al., 2012). Our model predicts that saturation of polarity factors at the OE enables the NE to compete more effectively for Cdc42 regulators and exceed the threshold of Cdc42 activation required for cell growth (Das et al., 2012).

Bifurcation analysis of the differential equations governing the model revealed that the progression from an asymmetric to a symmetric growth pattern travels through a landscape allowing symmetric and asymmetric states of active Cdc42 distribution at the cell tips, which are separated by a coexistence region, where both asymmetric and symmetric states are possible (Das et al., 2012). A prior model also predicted the role of coexistence and competition in fission yeast cell polarization; however, the effect was attributed to actin rather than Cdc42 (Csikasz-Nagy et al., 2008). Further, Cerone et al. (Cerone et al., 2012) described a similar coexistence region, without considering Cdc42 oscillations, as arising from increasing tip levels of the substrate for the polarisome, a Cdc42-regulated complex that consists of the polarity markers Tea1 and Tea4 and the formin For3. These authors proposed that the different morphological fates of *rga4*Δ sister cells, one of which remains monopolar and the other bipolar (Das et al., 2007), are due to each sister starting with a different initial condition, in a coexistence region that maintains itself through the cell cycle. This model implicated coexistence to explain the asymmetric morphological fates of *rga4*Δ cell (Das et al., 2007). More recent models of Cdc42 oscillations further explored the effects of diffusion through the cell and incorporated explicit molecular mechanisms for the positive feedback (Xu and Bressloff, 2016; Xu and Jilkine, 2018). All models are consistent with the idea that wild-type cells initially exhibit monopolar growth until the levels of Cdc42 regulators are sufficient to activate growth at the second cell tip. However, it is possible that environmental perturbations in wild-type cells or mutants that have defects in Cdc42 regulation may drive the system closer or further away from the coexistence region depending on the cell cycle stage, allowing alternative growth patterns to emerge.

To experimentally determine if different states of Cdc42 activation are possible in a biological context, and to identify novel molecular mechanisms that function to control morphological differentiation, we analyzed mutants that display alternative Cdc42 distributions in cells of the same size. Here, we describe the role of the Cdc42 GAP Rga4 in determining the initial state of Cdc42 GTPase activation in daughter cells. Our results suggest that the distribution of Cdc42 GAP proteins has crucial roles in determining the morphological fate of daughter cells and also ensuring consistent growth patterns from generation to generation.

## Results

### *rga4Δ* sister cells display divergent Cdc42 dynamics and distribution

We previously showed that the solutions of differential equations describing availability of Cdc42 regulators allow for a transition from asymmetric to symmetric states through a coexistence region, where different Cdc42 distributions at the cell tips are possible, leading to different patterns of cell growth (Supplemental Fig. 1A) (Das et al., 2012). This coexistence region is encountered as the cell grows and the availability of active Cdc42 or its regulators increases. In the model, the coexistence region arises when cells are around 7.6-8.0 micrometers long (Das et al., 2012). Since Cdc42 GTPase activation promotes bipolar cell growth when a threshold activity level has been achieved at the new cell end (Das et al., 2012), the timing of bipolar growth activation reflects the timing of the transition to a more symmetrical distribution of active Cdc42.

One prediction of the mathematical model is that the shape of the dynamical landscape as well as the initial state of the newly born cell specifies, within random fluctuations, the whole trajectory of Cdc42 dynamics as the cell grows (Supplemental Fig. 1A). Thus, we searched experimentally for molecular mechanisms that affect the coexistence region of Cdc42 activation and have a role in establishing the initial state of Cdc42 dynamics following cell division.

To this end, we searched for gene mutations that alter Cdc42 dynamics in newly born fission yeast daughter cells. We previously reported that a mutant strain lacking the Cdc42 GAP Rga4 displays asymmetric cell division, in which divergent growth patterns are observed in daughter cells (Das et al., 2012; Das et al., 2007). Thus, we examined the spatial distribution and activity of Cdc42 in *rga4*Δ daughter cells, as compared to *rga4+* control cells. We used the CRIB-GFP bioreporter (Tatebe et al., 2008) to visualize the localization and dynamics of the active form of Cdc42 (GTP-Cdc42) (Fig. 1A).

**Figure 1.**
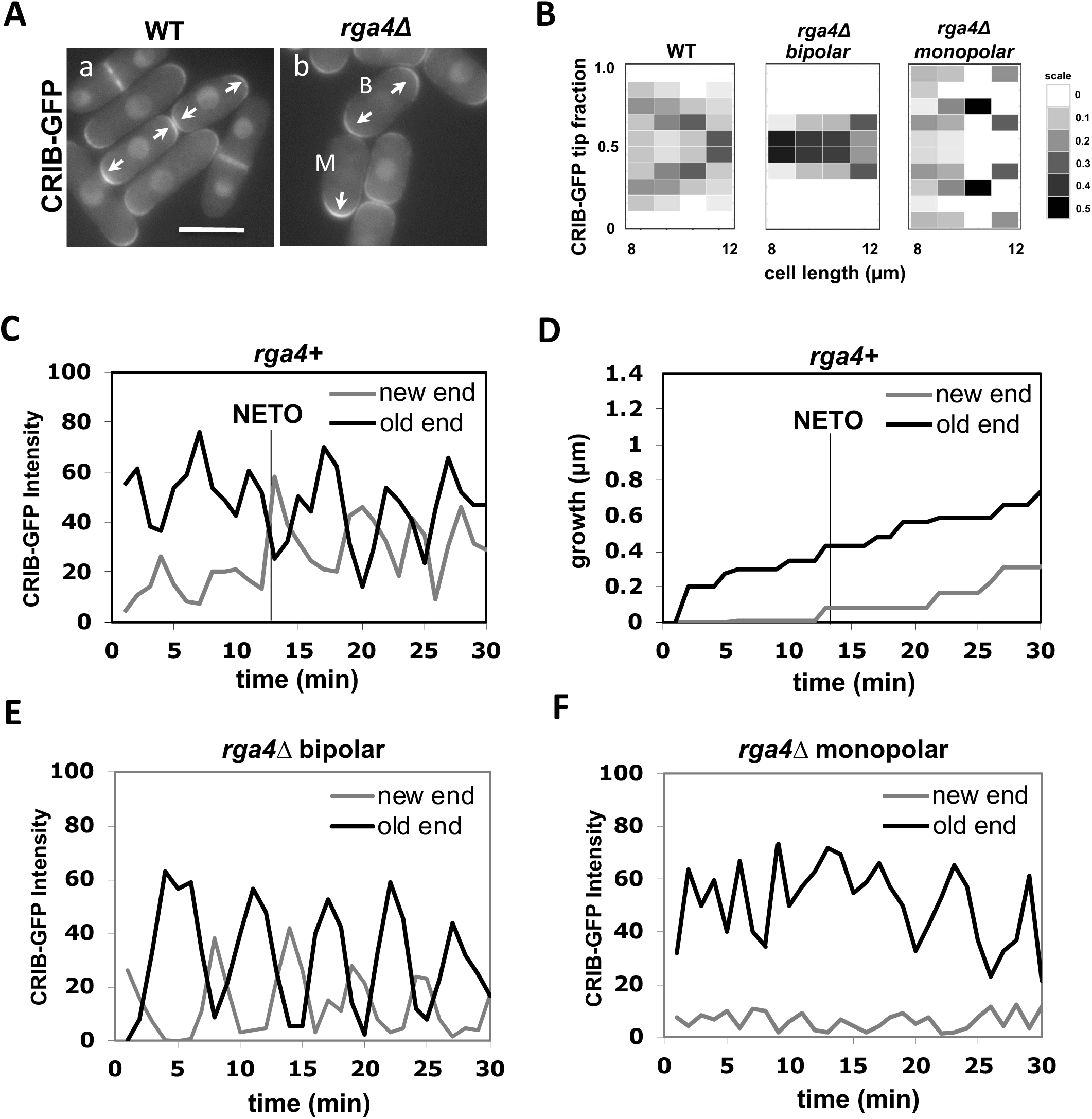
Loss of *rga4* leads to divergent patterns of Cdc42 dynamics. **A.** CRIB-GFP intensity in wild-type sister cells and *rga4*Δ sister cells. Scale Bar=5μm **B.** Heat map of CRIB-GFP distribution as a function of cell length in wild-type (N=61), *rga4*Δ bipolar (N=45), and *rga4*Δ monopolar (N=61) cells. **C.** CRIB-GFP intensity over time in the old end (black) and new end (gray) of a representative bipolar wild-type cell measured in 1-min intervals (N=5). **D.** Growth at the old (black) and new (gray) cell ends for the wild-type cell measured in C. The black line denotes the time at which the rate of growth changes in the new cell end (NETO). **E.** CRIB-GFP intensity over time in the old end (black) and new end (gray) of a representative bipolar *rga4Δ* cell (N=7) measured in 1-min intervals. **F.** CRIB-GFP intensity over time in the old end (black) and new end (gray) of a representative monopolar *rga4Δ* cell (N=7) measured in 1-min intervals.

As previously shown (Das et al., 2012), following cell division wild-type cells initially exhibit an asymmetric distribution of active Cdc42, where most of the active Cdc42 is localized at the old cell end (OE). As the cell increases in size, total levels of active Cdc42 increase, allowing Cdc42-GTP to accumulate at the new cell end (NE) resulting in a more symmetrical distribution of active Cdc42 (Fig. 1A, a; Fig. 1B). Both daughter cells display similar behaviors. Conversely, in cells lacking *rga4*, daughter cells exhibit different Cdc42 distributions after cell division, consistent with one daughter cell remaining monopolar and one daughter cell activating bipolar growth (Fig. 1A, b; Fig. 1B). In monopolar *rga4*Δ daughter cells, active Cdc42 remains more asymmetrically enriched at one cell end as the cell growths (Fig. 1A, b; cell “M”; Fig. 1B), whereas bipolar *rga4*Δ daughter cells display a more symmetrical distribution of Cdc42-GTP (Fig. 1A, b; cell “B”; Fig. 1B).

According to the model, active Cdc42 intensities oscillate at the cell tips due to a process of autoamplification, time-dependent negative feedbacks, and competition between cell tips. Wild-type cells initially display asymmetric oscillations of GTP-Cdc42, with most of the active Cdc42 at the old cell end. As the cell grows and levels of active Cdc42 increase, Cdc42-GTP oscillations become more symmetrical (Fig. 1C), allowing the cell to transition from monopolar to bipolar growth (Fig. 1D). Consistent with *rga4Δ* cells remaining competent for Cdc42 oscillations (Das et al., 2012), we found that *rga4Δ* daughter cells that activate bipolar growth display anticorrelated oscillatory dynamics of GTP-Cdc42 at both cell ends (Fig. 1E), while *rga4Δ* daughter cells that are destined to remain monopolar exhibit GTP-Cdc42 oscillations mainly at the OE (Fig. 1F). This variability indicates that the initial state of the cell may be different among *rga4Δ* daughter cells, allowing for more or less symmetrical distributions of active Cdc42.

### *rga4Δ* sister cells display divergent patterns of cell growth that are dependent on the growth pattern of the mother cell

Following cell division, wild-type daughter cells grow from the OE, the site that was previously growing in the mother cell (Fig. 2A, 2B). Activation of growth at the NE occurs later in the cell cycle, typically after the cell is longer than 9μm in length (NETO) (Fig. 1D) (Mitchison and Nurse, 1985). In contrast to wild-type cells, newly-born *rga4*Δ daughter cells exhibit divergent growth patterns (Das et al., 2007)(Fig. 2A). Some *rga4*Δ cells grow mainly from the OE, whereas other *rga4*Δ cells grow immediately form both ends, creating a population of *rga4*Δ mutant cells composed of both monopolar and bipolar cells (Fig. 2A) (Das et al., 2007).

**Figure 2.**
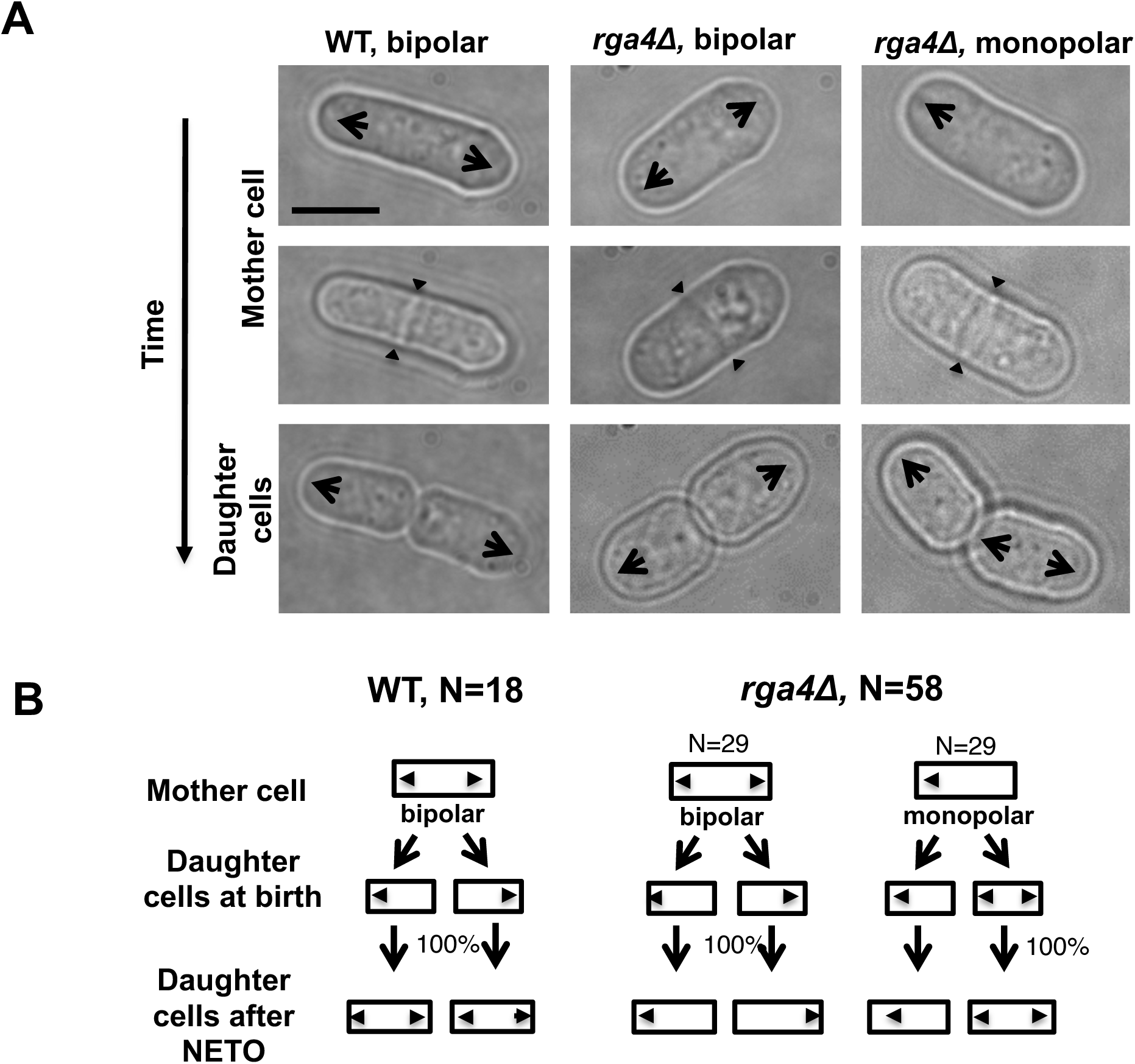
Loss of *rga4* leads to divergent patterns of growth. **A.** Time-lapse images showing initial growth patterns of daughter cells from bipolar wild-type, bipolar *rga4*Δ, and monopolar *rga4*Δ mother cells. Scale Bar=5μm. **B.** Diagram of the growth pattern of bipolar wild-type, monopolar *rga4Δ*, and bipolar *rga4Δ* mother cells and their respective daughter cells (showing the initial pattern of growth for daughter cells soon after birth).

To test if specific growth conditions in the mother cell affect polarized cell growth in the daughter cells, we performed a pedigree analysis in wild-type and *rga4*Δ cells by following cells in time by time-lapse microscopy. Wild-type mother cells generally grow in a bipolar fashion as they approach mitosis. Newly-born wild-type daughter cells first grow from the previously growing cell end (the “old” end), and then activate growth at the “new” end once they reach a minimal cell length (Mitchison and Nurse, 1985). This pattern of growth is similar in both daughter cells and is consistent from generation to generation (Fig. 2B). When we examined the growth patterns of *rga4Δ* daughter cells arising from monopolar or bipolar mother cells, we found that the specific pattern of growth of daughter cells depends on the history of the mother cells: *rga4Δ* monopolar mothers always give rise to one monopolar cell that inherited the previously growing tip and one bipolar cell that inherited a tip that did not previously grow (Fig. 2A, 2B). On the other hand, daughters of bipolar mother cells, which both inherited a previously growing tip, remain monopolar throughout the cell cycle. Thus, our observations indicate that the presence of a previously growing cell tip alters Cdc42 dynamics and cell growth in the daughter cell that inherits it.

### Mathematical modeling predicts different mechanisms of divergent Cdc42 dynamics in *rga4*Δ daughter cells

Since different distributions of active Cdc42 promote specific patterns of observed cell growth, we turned to mathematical modeling to explain the divergent Cdc42 dynamics observed in *rga4*Δ daughter cells. Our mathematical model predicts Cdc42 distribution and growth behavior of wild-type cells (Das et al., 2012)(Fig. 3A) and provides us with three different possible mechanisms to explain the growth pattern of *rga4*Δ cells (Fig. 3, B-D). For all three cases, we were led to assume an increase in the parameter C_sat_ describing the saturation activation threshold at each tip (compared to the parameter used for wild type cells; see Materials and Methods for details). This increase is anticipated in cells lacking GAP Rga4 that ought to be able to recruit larger concentrations of active Cdc42 and its regulators at the cell tip. The experimental observation of large amplitude oscillations for bipolar *rga4*Δ cells (Supplemental Fig. 1B) further indicates a stronger competition between tips (despite the larger saturation threshold), which leads us to assume an increase in either parameter ε describing the magnitude of the oscillation amplitude or parameter λ^+^_0_ describing the initial rate of Cdc42 activation at the lagging tip. These changes are consistent with a lack of Rga4 enabling stronger Cdc42 activation at both tips. Given these parameter changes, our model suggests three hypotheses that can be formulated mathematically (see Supplemental Materials for details):

1. The ability of each tip to activate Cdc42 depends on its prior growth history (Fig. 3B: Tip Prior Growth History) (Supplemental Fig. 2A). In the model of Fig. 3B, this was implemented by introducing a tip-aging parameter that describes the rate of Cdc42 recruitment at a given cell tip. This parameter is a function of the history of prior Cdc42 accumulation at the tip and it represents permanent features acquired by cell tips over long growth periods (such as stable membrane-associated macromolecular complexes or cell-wall mechanical properties). According to this model, daughters that inherit a tip that has experienced prior growth remain monopolar because the non-aged new tip cannot compete to accumulate Cdc42 and thus cannot age to the same extent as the dominant tip. By contrast, the daughter of a monopolar mother that inherits the tip that did not grow prior to division starts with two non-aged tips that compete equally and thus enter a bipolar oscillatory state in which they exchange Cdc42 and age together. According to hypothesis 1, the reason why wild type cells do not exhibit the *rga4*Δ growth pattern is that in wild type cells the reduced competition between the two cells tips (quantified in terms of different parameters for C_sat_, ε, λ^+^_0_) allows for NETO, after which both tips can grow and age prior to division (Fig. 3B, Supplemental Fig. 2A). Thus, following cell division, each wild-type daughter cell inherits similar Cdc42 histories, resulting in similar Cdc42 dynamics and growth patterns (Fig. 3B, Supplemental Fig. 2A).
2. Cell volume is a dominant factor in the *rga4*Δ growth pattern (Fig. 3C; Unequal Volumes of Daughter Cells). We assume that asymmetric cell growth of monopolar *rga4*Δ cells, persisting through the cell cycle to the point of cell division, could result in asymmetric cell shape or septum localization, generating daughters of unequal volume (but with approximately equal concentrations of Cdc42 and regulators otherwise). Because of the larger amount of Cdc42 and its regulators, the larger daughter could already be closer to the coexistence region, and thus start bipolar growth, exhibiting tip-to-tip oscillations, while the daughter with the smaller volume remains in the monopolar oscillatory region (Supplemental Fig. 1A). By contrast, daughters of bipolar mothers would divide in the middle, thus generating two daughters of equal volume, both growing in a monopolar fashion. Fig. 3C shows this is also a possible mechanism, though we note that this mechanism requires large volume differences between daughter cells (a 45% volume decrease for monopolar cells, and an 80% volume increase for bipolar cells, as compared to wild-type cells; Fig.3C), and may only partly explain the *rga4*Δ growth pattern: since wild-type cells go through the coexistence region in the process of doubling their volume, either avoiding (small cells), or directly entering (large cells) this region, the striking *rga4*Δ phenotype requires starting from very divergent daughter cell volumes, as compared to the initial volume of wild-type cells (Fig. 3C).
3. Unequal inheritance of Cdc42 regulators after division is the main contributor to the *rga4*Δ growth pattern (Fig. 3D; Unequal Distribution of Cdc42 Regulators) (Supplemental Fig. 2B). According to this hypothesis, the asymmetric growth of monopolar *rga4*Δ cells persists through the cell cycle to the point of cell division and thus the daughters of monopolar *rga4*Δ cells inherit different amounts of Cdc42 regulators (this is similar to hypothesis 2 except that the difference between the two daughters is concentrations instead of volume). One daughter could thus start and stay in the bipolar oscillatory region while the other remains in the monopolar region. By contrast, daughters of bipolar mothers generate two daughters with equal concentrations of regulators, both of which could grow in a monopolar fashion. This possible mechanism is illustrated in Fig. 3D, where we assumed that parameter λ^+^_0+_ (linear rate of Cdc42 activation) starts at different levels in the two daughters of monopolar mothers. In this model, cell growth results in the relaxation of concentrations towards a reference value, which thus allows the pattern to repeat itself (Supplemental Fig. 2B). Daughters of bipolar mothers maintain the reference concentrations inherited from their mother during growth (Supplemental Fig. 2B).

**Figure 3.**
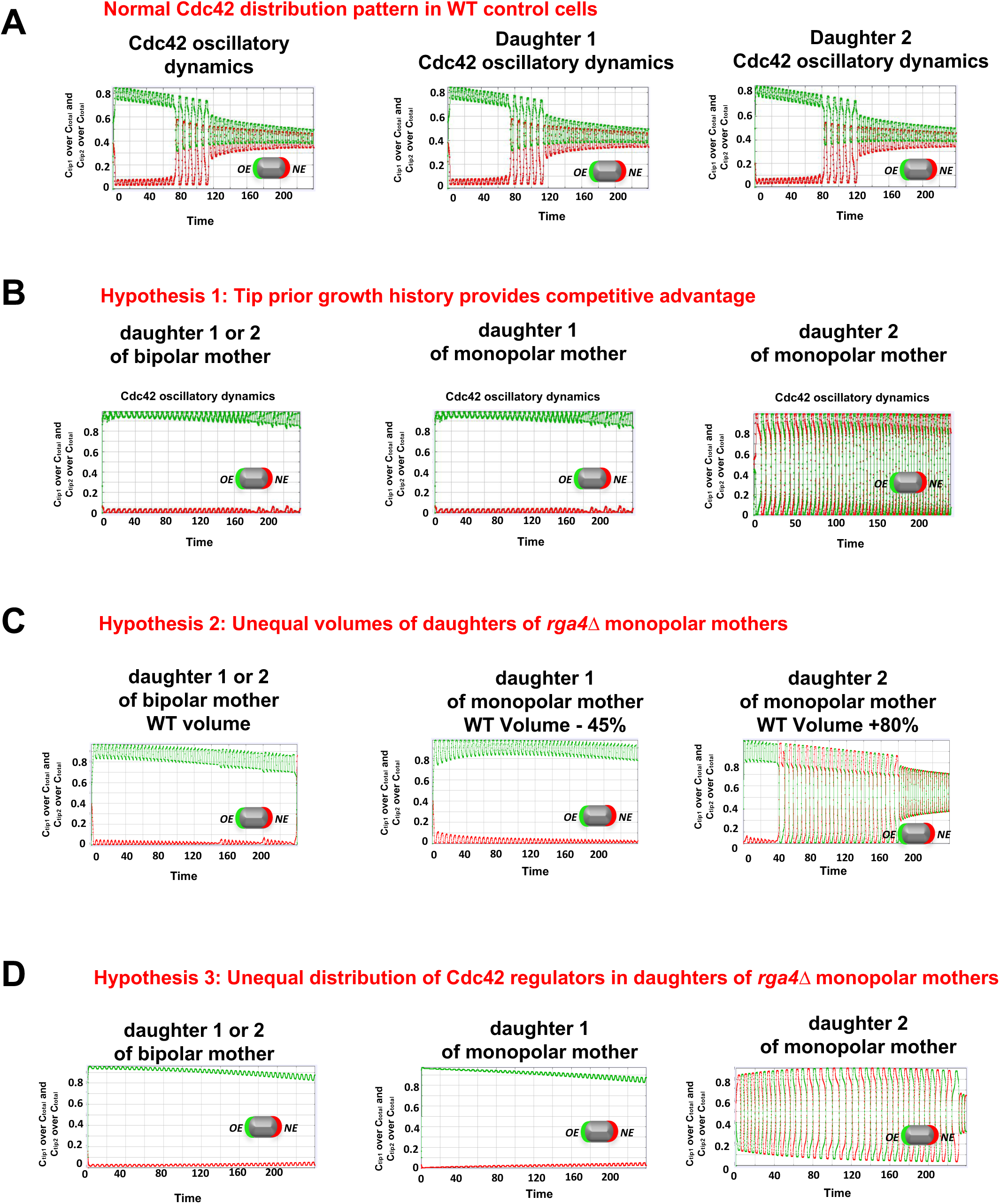
Model describing determinant of divergent Cdc42 dynamics in daughter cells. **A.** Model results for WT cells. The levels of active Cdc42 are indicated by a ratio between the tip bound Cdc42 over the total Cdc42 in the cell (C_tip_/C_total_). Plot shows WT control results of model with tip aging, which match the results of model by (Das et al. 2012) without tip aging; Cdc42 is shown at the old (green) and new (red) cell ends. Following cell division, both wild-type daughter cells inherit a Cdc42 history that results in cells having similar Cdc42 dynamics. The tip-aging parameter time is shown in Supplemental Fig. 2A. A difference to the model of (Das et al. 2012) without tip aging is that oscillations in the latter case are precisely symmetric, unlike the graphs in this panel in which the old end is stronger. **B.** Diagram showing the predicted effect of previous history of Cdc42 activation on the symmetry of Cdc42 activation in *rga4*Δ daughter cells (Hypothesis 1). The mathematical model that describes Cdc42 dynamics (Das *et al*, 2012) was modified by (1) increasing the saturation constant and oscillation amplitude and (2) incorporating a tip-aging parameter based on the history of Cdc42 activation at the cell tips. The tip-aging parameter versus time is shown in Supplemental Fig. 2A. **C.** Plots showing the effect of initial cell volume in *rga4*Δ cells (Hypothesis 2). Cells that start with half the volume of the mother or less (first two graphs) remain asymmetric while those that start with larger volume (third graph) encounter the coexistence region and exhibit symmetric oscillations for most part of their growth, corresponding to bipolar growth. **D.** Results of model with unequal distribution of Cdc42 regulators, represented by a growth-dependent rate constant (Hypothesis 3). The corresponding dependence of 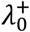 on time is shown in Supplemental Fig. 2B.

The above three hypotheses for the *rga4*Δ polarization pattern correspond to three ways in which the daughter cells of monopolar mothers are physically different to each other. A final fourth possibility is that unlike in Hypotheses 1-3, these sister cells have in fact the same physical properties (volumes, concentrations, *etc.*) but they end up growing differently because of the initial conditions of Cdc42 distribution: they happen to start in different basins of attraction of a co-existence region that spans the whole cell cycle (Cerone et al., 2012).

We performed a series of experiments to test these hypotheses, using genetic screening methods to identify factors that alter the growth patterns and Cdc42 distribution in *rga4*Δ daughter cells.

### A genome-wide screen identified determinants of Cdc42 dynamics in *rga4*Δ daughter cells

In order to identify polarity regulators that may functionally interact with Rga4, we performed a synthetic genetic array (SGA) screen to identify mutations that suppress or enhance the polarity and growth pattern defects observed in *rga4Δ* mutants. We crossed the *rga4Δ* strain (FV1529) with the Bioneer fission yeast deletion library (Kim et al., 2010) and visually screened the double mutants for morphology and colony growth phenotype. This analysis identified several regulators of polarity and cytokinesis as functionally interacting with Rga4 (Fig. 4). Gene functions that normalize the growth pattern of *rga4*Δ mutants, when deleted, include a regulator of Cdc42 activation, the Cdc42 GEF Gef1 (Das et al., 2015), and a factor involved in division site placement, the DYRK family protein kinase Pom1 (Tatebe et al., 2008). Conversely, loss of other Cdc42 regulators enhanced the morphological defects of *rga4*Δ mutants; these include the Cdc42-dependent Pak1/Shk1 kinase (also known as Orb2) (Marcus et al., 1995), which is involved in negative-feedback control of Cdc42, the Cdc42 GDI Rdi1 (Bendezu et al., 2015), the Cdc42 GAP Rga6 (Revilla-Guarinos et al., 2016) and PP2A activator, Ypa2/Pta2 (Bernal et al., 2012). Loss of *efc25*, a Ras1 GEF that controls Ras1 GTPase activity and promotes Cdc42 polarization, produced round cells in the *rga4*Δ background. The functional interaction of Rga4 with Ypa2/Pta2 and with Cdc42 regulator Rga6 have been previously reported (Bernal et al., 2012; Revilla-Guarinos et al., 2016). Thus, gene functions that normalize the pattern of growth of the bipolar daughter cell, preventing precocious bipolar growth activation, decrease Cdc42 activity (*gef1*) or affect the placement of the site of division (*pom1*), while genes that enhance the morphological asymmetry of *rga4*Δ cells encode negative Cdc42 regulators (*rga6, pak1/shk1*).

**Figure 4.**
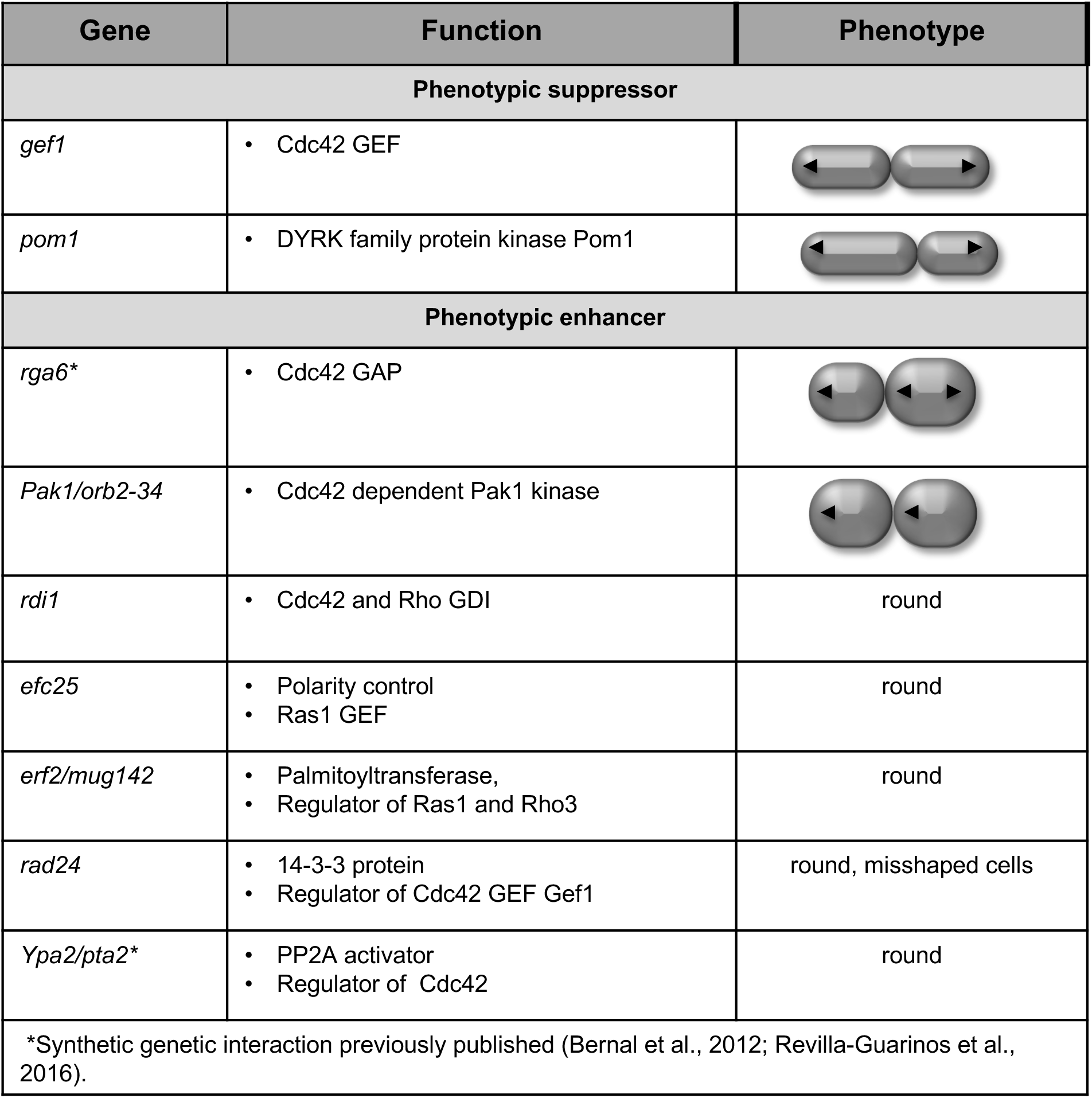
Rga4 interacts with various polarity factors affecting the initial growth patterns and/or morphology of *rga4*Δ daughter cells. Synthetic genetic array screen to identify factors that alter the growth pattern of *rga4*Δ cells. Loss of the Cdc42 GEF Gef1 and the DRCK kinase Pom1 suppress aspects of the *rga4*Δ phenotype. Loss of the Cdc42 negative regulators, the guanine dissociation inhibitor Rdi1 and the Cdc42 GAP Rga6 worsen the morphology of *rga4*Δ bipolar cells. A mutant allele of the essential Cdc42 dependent Pak1 kinase (*orb2-34*) is viable in combination with *rga4*Δ at 25°C (with an altered morphology) but is synthetic lethal with *rga4*Δ at 36°C. The functional genetic interactions of *rga4* with *rga6* and *ypa2/pta2* have been previously published (Bernal et al., 2012; Revilla-Guarinos et al., 2016).

### Localization of the Cdc42 scaffold Scd2 is not altered in *rga4*Δ daughter cells

The asymmetric Cdc42 dynamics and growth patterns observed in *rga4Δ* daughter cells can be explained by the presence of a “tip aging” factor, that accumulates as each tip grows (Hypothesis 1 in Fig. 3B). An attractive candidate for this function is the scaffold Scd2, a component of the polarisome, that has an important role in Cdc42 autoamplification (Lamas et al., 2020). Scd2 average distribution correlates with Cdc42 levels and cell growth at the tips: during interphase, Scd2-GFP localizes asymmetrically in monopolar wild-type cells, whereas in bipolar wild-type cells Scd2-GFP is distributed more symmetrically (Fig. 5A; Supplemental Fig. 3A). Conversely, in *rga4*Δ daughter cells, Scd2-GFP distribution is very symmetrical even in small bipolar cells, while it is mostly asymmetrical in monopolar *rga4*Δ cells (Supplemental Fig. 3A).

**Figure 5.**
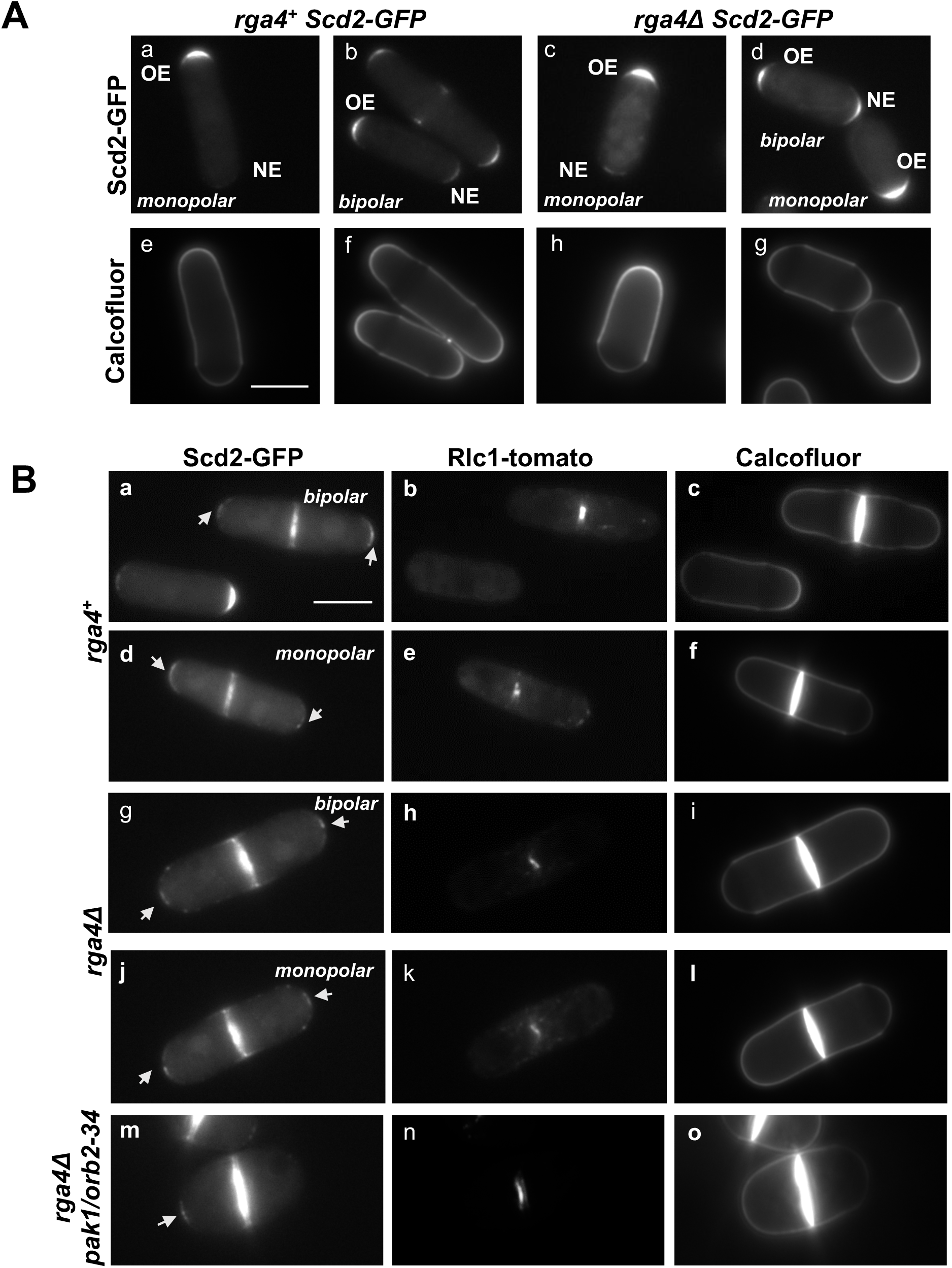
Tip localization of small amounts Scd2-GFP remains at both tips during cell division in wild-type and *rga4*Δ cells. **A.** Scd2-localization in wild-type and mutant *rga4*Δ monopolar and bipolar cells. **B.** Images of Scd2-GFP (arrows) in septated cells during constriction of the actomyosin ring, visualized using Rlc1-tomato. Scd2-GFP is present at the tips in wild-type, *rga4Δ*, and *rga4Δ pak1/orb2-34* mother cells during actomyosin ring constriction.

Thus, to test if Scd2 has a role as a “tip aging” factor, that “marks” the previously growing tip, we analyzed the distribution of the Cd42 scaffold Scd2 during cell division in wild-type and *rga4*Δ cells. Remarkably, we observed that a small amount of Scd2 remains localized to the cell tips in wild-type cells undergoing mitosis and cytokinesis (Fig. 5B, a-f), albeit at a much lower level as compared to interphase cells (Fig. 5A). Both bipolar and the few monopolar wild-type cells display Scd2 markings at the tips. This novel finding indicates that Scd2 may indeed serve as a tip marker, conferring memory of the history of Cdc42 activation at previously growing cell tips.

Thus, we hypothesized that monopolar *rga4*Δ cells may exhibit asymmetric localization of Scd2-GFP between previously growing and non-growing tips, and that this imbalance may account for the different growth patterns and Cdc42 dynamics in *rga4*Δ daughter cells. When we analyzed monopolar and bipolar mother cells, however, we found no significant difference in the levels of Scd2-GFP during cell division at the two tips in monopolar and bipolar cells from both wild-type and *rga4Δ* mutant cell cultures (Fig. 5B, g-l; Supplemental Fig. 3B).

To test if inheritance of polarisome components has an effect on cell growth pattern in daughter cells, we altered Scd2 localization in *rga4*Δ cells by introducing the *pak1/orb2-34* mutation, which leads to Scd2 accumulation only at one tip (Das et al., 2012). These *rga4*Δ *orb2-34* mutants grow only in a monopolar manner (Fig. 4). We found that Scd2 localizes only to one tip during mitosis in *rga4*Δ *orb2-34* mutants, the end that grew previously (Fig. 5B, m-o). In this case, the usually bipolar *rga4*Δ daughter cell which does not inherit any growing ends, defaults to activate cell growth only from the area of the septum where Scd2 is located (Fig. 5B, m; Fig. 4).

These results suggest that, while Scd2 permanence at the cell tips may have a role in “marking” growing cell tips and promoting cell polarization after mitosis, the divergent Cdc42 dynamics observed in *rga4Δ* sister cells do not arise from asymmetric inheritance of the Scd2 complex at the cell tips.

### Decreasing the initial cell size of bipolar *rga4*Δ daughter cells abolishes bipolar growth

Because an increased cell volume could explain the symmetrical Cdc42 distribution of *rga4* bipolar cells, we tested if reducing the size of the daughter cell destined to be bipolar could normalize its pattern of growth. Interestingly, we identified the kinase Pom1 as a suppressor of the *rga4*Δ phenotype. The DYRK kinase Pom1 is an established regulator of the placement of the division site (Bahler et al., 1998; Rincon et al., 2014). In cells that lack *pom1*, the cell septum forms closer to the new cell end, generating daughter cells with different volumes (Bahler et al., 1998). Loss of *pom1* in *rga4*Δ mutant cells indeed shifted the site of cell division towards the new cell end, particularly in monopolar *rga4*Δ mother cells (Fig. 6A). This shift decreased the initial volume of the daughter cell inheriting the non-growing new end, which would normally be destined to be bipolar (Fig. 6C). When we followed the growth pattern of daughter cells, we found that all monopolar *pom1*Δ *rga4*Δ mother cells gave rise to two monopolar daughter cells (Fig. 6A) (n=10), whereas all monopolar control *rga4*Δ mother cells gave rise to one monopolar and one bipolar cell (n=18). These observations suggest that decreasing the initial size of the cell that inherits a non-growing tip prevents precocious bipolar growth activation. Interestingly, the increased size of the cell that inherits the growing tip did not cause it to switch to bipolar growth.

**Figure 6.**
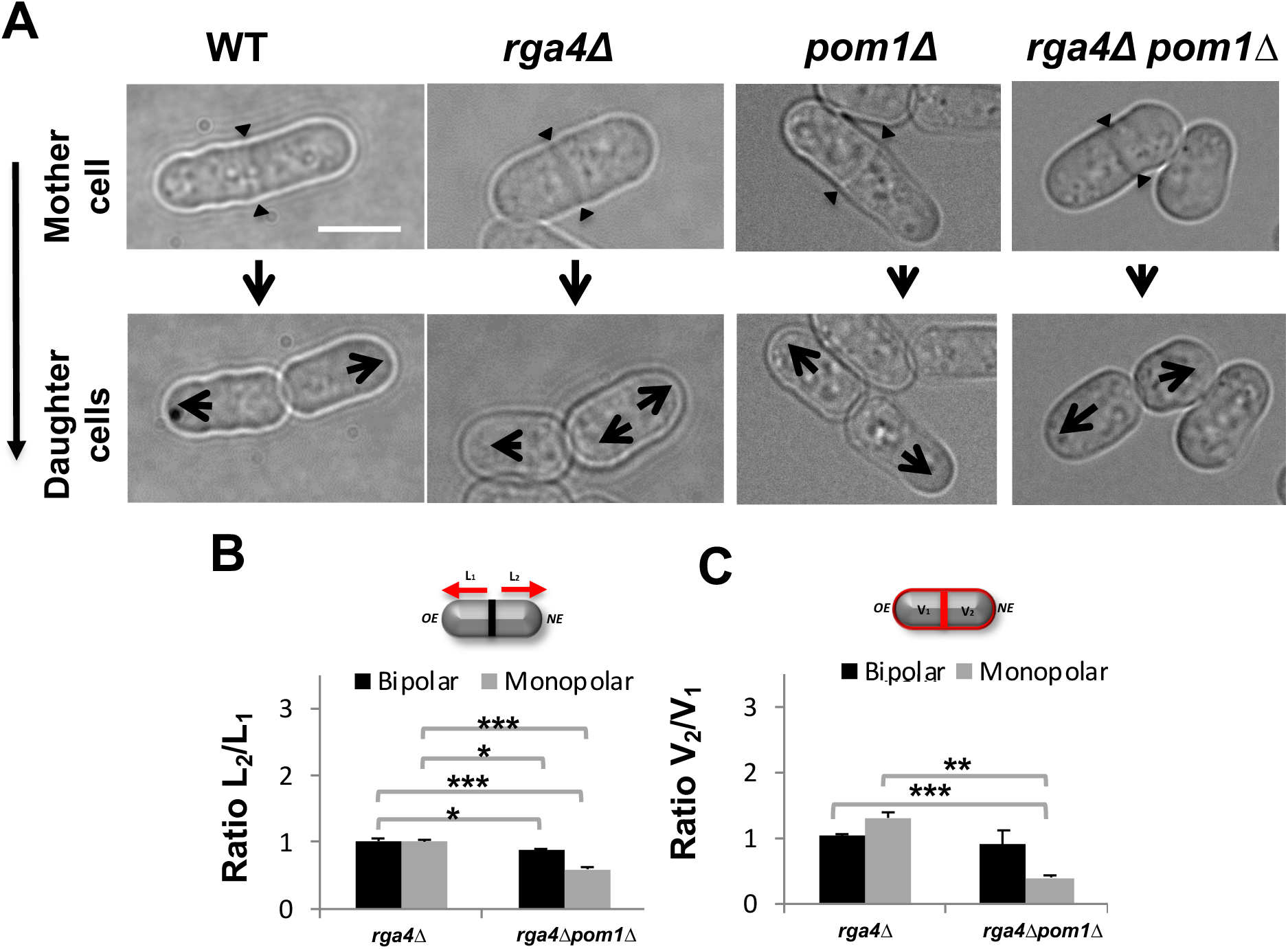
Loss of DYRK-family kinase Pom1 suppresses precocious bipolar cell growth in *rga4* mutant cells. **A.** Pattern of growth in WT, *rga4*Δ and *rga4*Δ *pom1*Δ cells. Scale bar=5μm. Large arrows indicate direction of mono- or bi-polar growth. Small triangular arrows indicate sites of septation. **B.** Quantification of the ratio of the distances between the septum to the cell tip in septated *rga4*Δ (bipolar N=10, monopolar N=10) and *pom1*Δ *rga4*Δ (bipolar N=5, monopolar N=19) cells. Statistical analysis was done using ANOVA followed by Tukey HSD post hoc test (p=0.011; p=000; p=0.018; p=000) based on 3 independent experiments. **C.** The quantification of the ratio of the volume in the two compartments in septated cells. Statistical analysis was done using ANOVA followed by a Tamhane post hoc test based on 3 independent experiments (p=0.000; p=0.006).

### Cdc42 GTPase regulators control the cell growth pattern of *rga4*Δ daughter cells

The presence of different amounts of Cdc42 regulators can also account for divergent Cdc42 dynamics in *rga4*Δ daughter cells. SGA analysis revealed a suppressor interaction between Rga4 and Cdc42 GEF Gef1 (Fig. 4). We have previously shown that Gef1 and Rga4 have opposing effects in the control of Cdc42 activation at the cell tips and cell diameter (Das et al., 2015). Cells expressing a mutated form of *gef1* (*gef1S112A)*, that increases Gef1 cortical localization, exhibit precocious bipolar growth activation (Das et al., 2015). Thus, we tested if loss of *gef1* suppresses the distinct growth pattern of *rga4*Δ daughter cells by reducing the activity of Cdc42 in the bipolar cell. Indeed, we found that cell growth in *rga4*Δ *gef1*Δ daughter cells was normalized, with a significantly increased percentage of daughter cells from monopolar *rga4*Δ mother cells growing in a monopolar manner, as compared with control *gef1+ rga4*Δ cells (Fig. 7A, 7B, 7C). Hence, decreased levels of Cdc42 suppresses the precocious cell growth at the NE of *rga4*Δ daughter cells destined to be bipolar.

**Figure 7.**
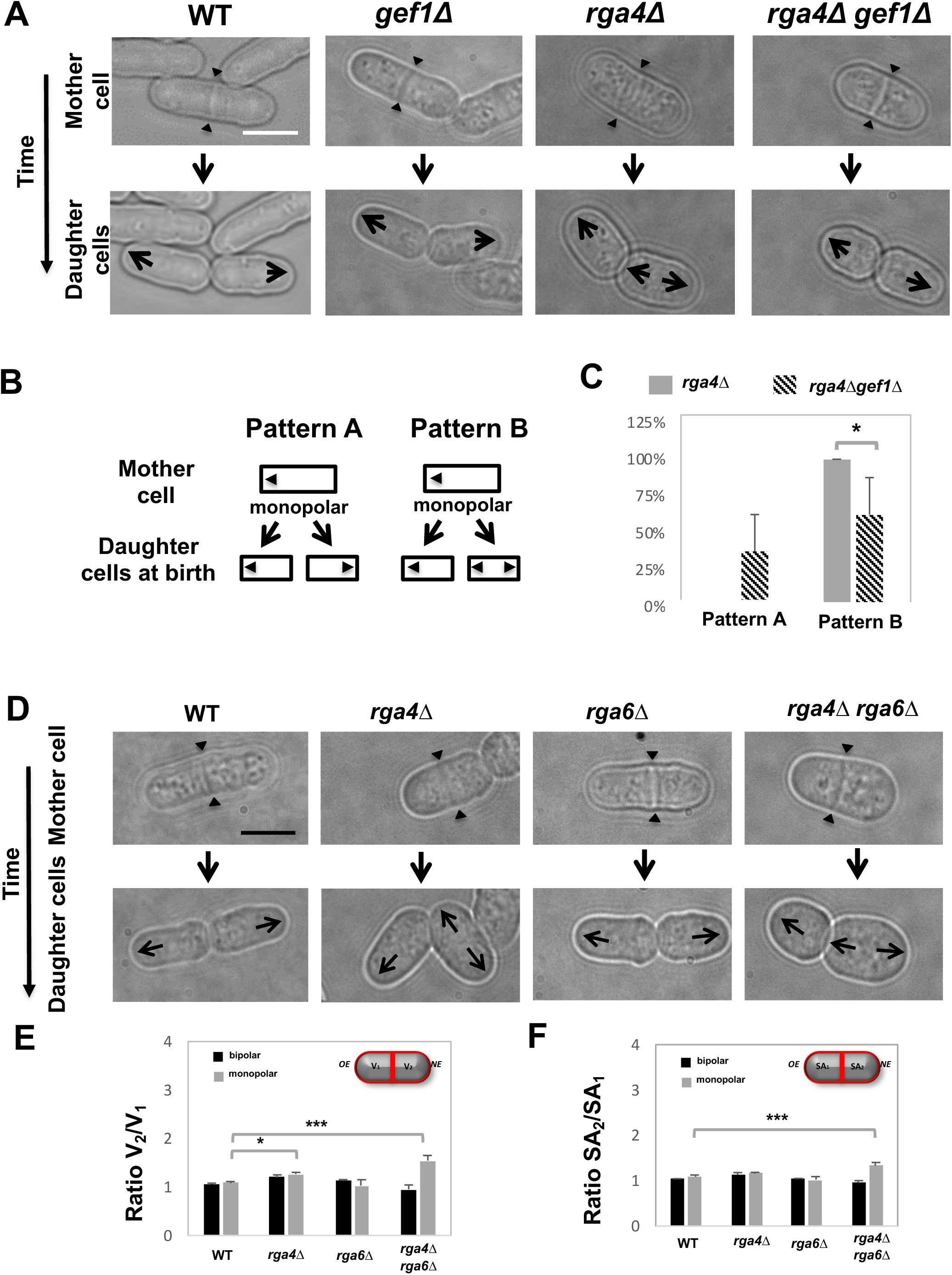
**A-C. Loss of Cdc42 GEF Gef1 increases the percentage of monopolar mother cells that produce daughter cells with similar growth patterns. A.** Time-lapse DIC images of wild-type, monopolar *gef1*Δ, monopolar *rga4*Δ and monopolar *rga4*Δ *gef1*Δ cells showing the growth patterns of daughter cells. **B.** Diagram showing two alternative growth pattern of daughter cells born from monopolar cells. In pattern A, both daughter cells grown in a monopolar fashion after birth; whereas in pattern B, one daughter cell growths in a monopolar fashion and the other daughter cells exhibits bipolar growth after birth. **C.** Quantification of the proportion of *rga4*Δ (N=29) and *rga4*Δ *gef1*Δ (N=14) monopolar mother cells exhibiting either growth pattern A (two monopolar daughter cells) or growth pattern B (one monopolar and one bipolar daughter cell). Statistical analysis was done using an independent T-test with SPSS statistics package 22.0 based on 6 independent experiments (p=0.013). **D-F. Loss of *rga4*Δ and *rga6*Δ produces daughter cells with different cell volumes. D.** Time-lapse DIC images of wild-type, *rga4*Δ, *rga6*Δ, and *rga4*Δ *rga6*Δ cells undergoing cell division. Arrow heads indicate the position of the cell septum. Scale bar=5μm. **E.** Quantification of the ratio of the cell volumes of the two compartments in septated cells in wild-type (bipolar N=63, monopolar N=12), *rga4*Δ (bipolar N=52, monopolar N=17), *rga6*Δ (bipolar N=58, monopolar N=5), and *rga4*Δ *rga6*Δ (bipolar N=21, monopolar N=43) strains. Statistical analysis was done using analysis of variance (ANOVA) with SPSS statistics package 22.0, followed by Tukey HSD post hoc test based on 3 independent experiments (p= 0.018; p=0.000; p=018; p=0.000). **F.** Quantification of the ratio of the surface area in the two compartments as previously done in B. Statistical analysis was done using ANOVA followed by Tukey HSD post hoc test based on 3 independent experiments (p= 0.022 p=0.000)

SGA analysis also revealed a functional interaction between Rga4 and another Cdc42 GAP in *S. pombe*, Rga6 (Revilla-Guarinos et al., 2016) (Fig. 4). We found that, consistent with the partially overlapping functions of Rga4 and Rga6 (Revilla-Guarinos et al., 2016), loss of *rga6* in the *rga4*Δ mutant exacerbates the asymmetric morphology observed in *rga4*Δ daughter cells (Fig. 7D). Using the Pombe Measurer ImageJ plugin (F. Chang, Columbia University), we calculated the volume (Fig. 7E) and surface area (Fig. 7F), of the two cell compartments in septated monopolar and bipolar mother cells from WT, *rga4*Δ and *rga4*Δ *rga6*Δ cell cultures. We found that loss of Rga6 significantly increased the volume (V_2_/V_1_) (Fig. 7E) and surface area (SA_2_/SA_1_) (Fig. 7F) ratios of the two compartments in the *rga4*Δ background, as compared to wild-type cells (where V_1_ or SA_1_ are the OE compartment and V_2_ or SA_2_ are the NE compartment). These results suggest that daughter cells from monopolar mothers inherit divergent starting cell morphologies and cell volumes: the *rga4*Δ cell inheriting the previously growing end is generally slightly smaller, a phenotype exacerbated by loss of *rga6* (Fig. 7D). Thus, loss of the second Cdc42 GAP, Rga6, reveals that the “bipolar” daughter cell that inherits the NE has different morphological properties than the “monopolar” daughter cell that inherits the OE.

### Cdc42 GAP Rga6 distribution is asymmetrically inherited during cell division in *rga4*Δ cells

Since loss of Cdc42 GAP Rga6 exacerbated the asymmetric phenotype of *rga4*Δ mutants, we investigated whether loss of Rga4 alters the distribution of Rga6 at the cell membrane. To test this, we measured the intensity of Rga6-3YFP throughout the cell membrane in interphase wildtype and *rga4*Δ cells. We found that monopolar *rga4*Δ cells exhibit an asymmetric distribution of Rga6-3YFP which is increased near the OE and depleted from the NE of the cell (Fig. 8A, b). Conversely, wild-type cells (Fig. 8A, a) and bipolar *rga4*Δ cells (Fig. 8A, c) display a more symmetrical Rga6 distribution, enriched near both the OE and NE. Importantly, we found that this distribution is maintained throughout mitosis, and is inherited by the daughter cells (Fig. 8B and Fig. 8C). In dividing monopolar *rga4*Δ cells, the daughter cell that inherits the growing end (OE cell, Fig. 8C) also acquires increased amounts of Rga6 (Fig. 8B, a,b), as compared to the daughter cells that inherits the non-growing end (NE cell). Conversely, this asymmetric inheritance is not found in wild-type cells (Fig. 8C) or in *rga4*Δ bipolar cells (Fig. 8C, Fig. 8B, c and d). Rga6 localization is likely determined by the specific pattern of growth of cells during interphase (Fig. 8A), where we find Rga6 localization to be more asymmetrical early in the cell cycle in monopolar cells, to become more symmetrically distributed once NETO occurs.

**Figure 8.**
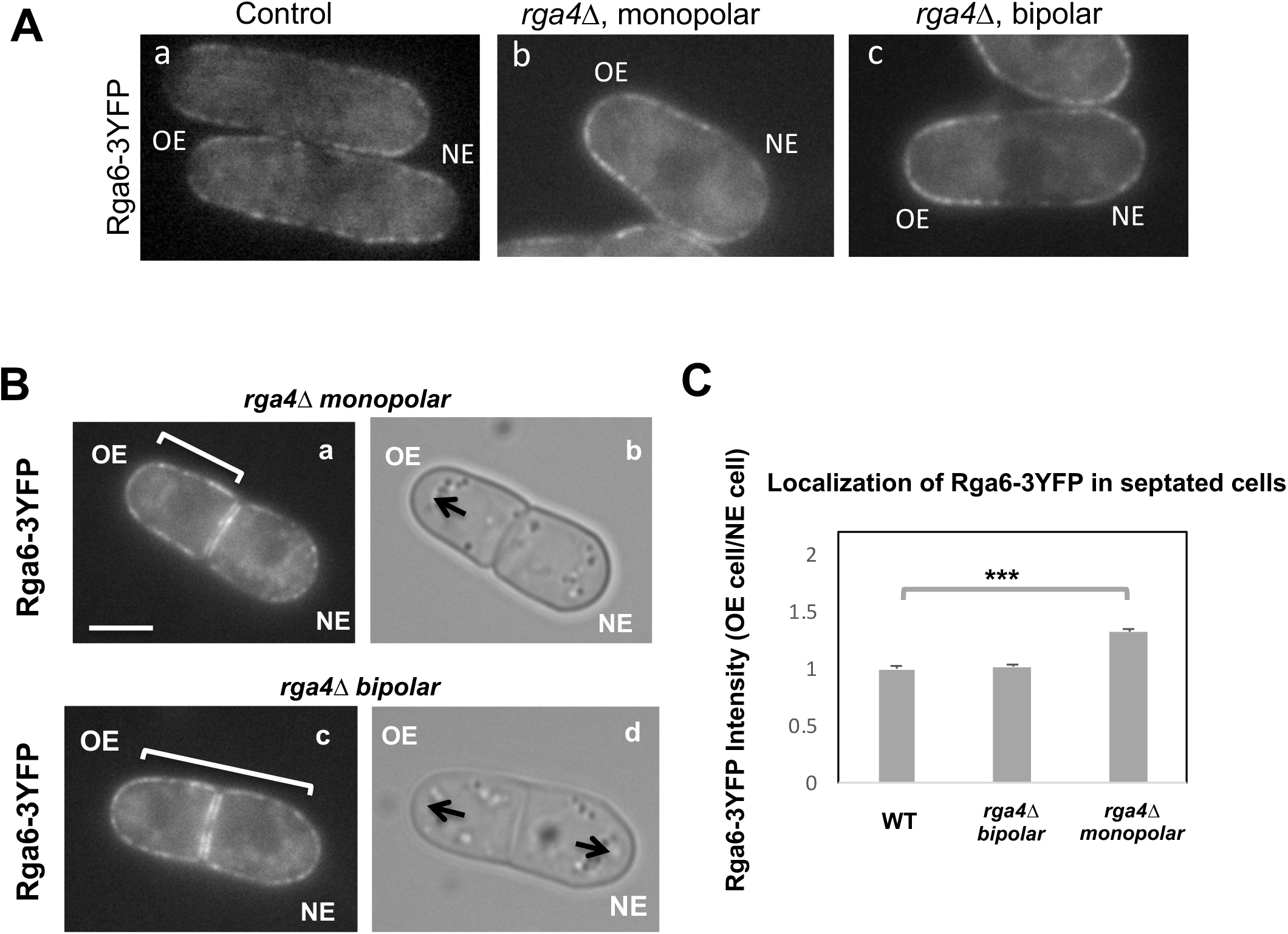
Distribution of Rga6 to daughter cells is altered in monopolar or bipolar *rga4*Δ monopolar mother cells. **A.** Distribution of Rga6-3YFP in interphase control, monopolar *rga4*Δ and bipolar *rga4*Δ cells. **B.** Localization of Rga6-3YFP in *rga4*Δ dividing cells. Scale bar=5 μM. **C.** Ratio of Rga6-3YFP intensities at the cell membrane, in the OE containing compartment over the NE containing compartment, in septated cells; in bipolar wild-type (N=25), *rga4*Δ bipolar (N=25), and *rga4*Δ monopolar (N=31) cells. Statistical analysis was done using ANOVA followed by Tukey HSD post hoc test based on 3 independent experiments (p= 0.000). **C.** Distribution of Rga6 in interphase control, monopolar *rga4*Δ and bipolar *rga4*Δ cells.

The localization of two other Cdc42 regulators, the guanine nucleotide exchange factors (GEFs) Scd1 and Gef1, do not display visible differences in dividing monopolar or bipolar *rga4*Δ mother cells (Supplemental Fig. 4). The third Cdc42 GAP, Rga3 (Gallo Castro and Martin, 2018), displays a very faint localization at the cell tips of dividing cells (Supplemental Fig. 4), which did not seem significantly more asymmetrical in dividing *rga4*Δ monopolar mother cells (Supplemental Fig. 4).

These observations, showing that daughter cells of *rga4*Δ monopolar cells inherit different amounts of Cdc42 GAP Rga6 at the membrane may explain how different distributions of active Cdc42-GTP are present in *rga4*Δ sister cells immediately following cytokinesis. Taken together, our results support the notion that divergent *rga4*Δ daughter cells are born with distinct initial conditions, resulting in different growth patterns. We show that negative regulators of Cdc42 activation, the Cdc42 GAPs Rga4 and Rga6, have a crucial role in ensuring that *S. pombe* daughter cells follow similar morphological destinies. Since Rga6 localization becomes more biased towards the growing cell end in monopolar *rga4*Δ cells, these findings suggest that Cdc42 GAP proteins provide a link between the history of cell growth in the mother cell and the initial state of Cdc42 activation in daughter cells.

## Discussion

### The pattern of growth in the mother cell determines the Cdc42 oscillatory dynamics and distribution in daughter cells

We previously showed that Cdc42 exhibits oscillatory dynamics, and that the relative amounts of active Cdc42 at the cell tips determines the pattern, either monopolar of bipolar, of cell growth (Das et al., 2012). We proposed a mathematical model that describes the distribution of active Cdc42 GTPase at the cell tips in fission yeast cells (Das et al., 2012), and one of the predictions of this model is that the initial state of Cdc42 distribution in newly born daughter cells has an important role in defining the future behavior of the system. Thus, in this paper, we asked the question: what determines the initial state of Cdc42 dynamics in newly born daughter cells? To answer this question, we analyzed Cdc42 dynamics in *rga4*Δ cells, that are missing one of the three fission yeast Cdc42 GTPase activating proteins, Rga4. We previously reported that loss of *rga4* confers divergent patterns of growth in daughter cells, where one cell remains monopolar, while the other displays precocious activation of bipolar growth (Das et al., 2007). Here, we show that, consistent with their mode of growth, *rga4*Δ daughter cells show divergent Cdc42 dynamics: one daughter displays highly asymmetric, while the other highly symmetric, active Cdc42 dynamics and distribution. Further, we find that the particular Cdc42 dynamics of *rga4*Δ cells are determined by the mode of growth, either monopolar or bipolar, of the mother cell. Thus, these observations reveal a link between the mode of cell growth in the previous cell cycle and the morphological fate of daughter cells.

In wild-type cells, cells start to grow from the “New” end, and thereby activate bipolar growth, in a process called “NETO” (Mitchison and Nurse, 1985) usually later in the cell cycle, during the G2 phase. Thus, once the mother cell divides, both daughter cells inherit one end that had previously been growing (the “old” ends), and following cell division both daughter cells behave similarly, activating the OEs first. The functional purpose of this process has been so far been relatively elusive. In *rga4Δ* cells, the consequences of not activating NETO become more evident: if the mother cell is monopolar, the daughter cell that inherits the never-activated cell tip displays symmetric Cdc42 activation, and grows symmetrically from both cell ends, generating a different morphology than its sister cell. Our observations suggest a novel function for NETO, in enabling cells to maintain consistent Cdc42 dynamics and cell polarization, across generations.

### A mathematical model of Cdc42 oscillations predicts different mechanisms for divergent Cdc42 dynamics in daughter cells

The distinct oscillatory dynamics of Cdc42 in monopolar and bipolar *rga4Δ* cells can be explained by variations of our mathematical model (Das et al., 2012). The model predicts three different ways that reproduce the Cdc42 dynamics observed in the *rga4*Δ phenotype.

The first way invokes a tip-aging parameter that produces an asymmetry in the Cdc42 history between the two cell tips. In this hypothesis, the divergent behavior of *rga4*Δ sister cells results from inheriting different levels of an “end marker” that accumulates with cell growth. Mathematically, this model was implemented by introducing tip-aging dimensionless parameters *a*_1_ and *a*_2_, which describe how the rate of Cdc42 recruitment at either tip increases as a result of a history of prior growth. Following cell division, active Cdc42 is predicted to oscillate around different averages at the two cell ends, as we observe in monopolar *rga4*Δ cells, that have inherited one “marked” end. Conversely, the lack of previous Cdc42 activation at both cell tips, and thus a more symmetric Cdc42 history, yields similar average Cdc42 activation at the two cell ends and more symmetric Cdc42 oscillations, as we observe in bipolar *rga4Δ* cells (Das et al., 2012).

The second hypothesis involves asymmetric cell division, where the daughter cells inherit different cell volumes. Cdc42 dynamics are affected by the concentrations of Cdc42 regulators available to activate Cdc42. Larger cell volumes, increasing the availability of cytoplasmic Cdc42 regulators, lead to increased Cdc42 activity at the new end and thus to more symmetrical Cdc42 oscillations at both ends (Das et al., 2012). Mathematically, daughters of monopolar *rga4*Δ mothers cells start with different total amounts *C*_*tot*_, in proportion to the differences in volume. The equations for this model are the same as the ones in (Das et al., 2012).

Finally, the third hypothesis invokes the asymmetric inheritance of Cdc42 regulators, either GEFs (Guanine nucleotide Exchange Factors) or GAPs (GTPase Activating Proteins) in the two daughter cells. Mathematically, parameter λ^+^_0_ (linear rate of Cdc42 activation) starts at different levels in the two daughters of monopolar mothers; cell growth results in the relaxation of concentrations represented by λ^+^_0_ towards a reference value. This effect would lead to different Cdc42 dynamics in the two daughter cells: indeed, increasing the level or availability of the Cdc42 activator Gef1 by overexpression, or by specific mutations, leads to more symmetrical Cdc42 oscillations at both ends (Das et al., 2012; Das et al., 2015).

### Mutant alleles of Cdc42 regulators functionally interact with *rga4*, modulating Cdc42 dynamics and growth pattern in *rga4*Δ daughter cells

To distinguish between the different hypothesis of the model, we used a Synthetic Genetic Array (SGA) screen to search for gene functions that modulate the pattern of cell growth in *rga4*Δ daughter cells. We identified several suppressors that drive the bipolar daughter cell to grow in a monopolar manner when deleted in the *rga4*Δ background, including *gef1* (encoding a Cdc42 GEF (Coll et al., 2003)), and *pom1* (encoding a DYRK kinase that controls the placement of the septum (Bahler and Nurse, 2001). Our SGA screen also revealed functional interactions that enhanced the morphological phenotypes of *rga4*Δ cells: in particular, other negative regulators of Cdc42, including the Cdc42 dependent Pak1 kinase Pak1/Shk1/Orb2-34, the Cdc42 GDI Rdi1, and the Cdc42 GAP Rga6. Overall, this screen indicated that precocious bipolar growth and shape asymmetry of *rga4*Δ daughter cells could be ameliorated by decreasing Cdc42 activation and/or decreasing the cell volume of the cell inheriting the never-grown tip; and it could be enhanced by loss of the second Cdc42 GAP, Rga6.

### Genetic analysis to test the different hypotheses of the model

To test the three alternative model hypotheses, we followed the localization of Cdc42 complex components, or we changed the overall cell behavior by introducing gene mutations that alter Cdc42 activity.

Hypothesis 1 invokes the presence of a tip-aging parameter. When introduced into our mathematical model, this tip-aging parameter models how cell growth progressively “marks” the cell tip in the mother cell. The “mark” accumulates at higher values at the tips which grow for the longest time (OE) and then remains at the cell tips throughout mitosis. In the following generation, the “mark” promotes the activation of Cdc42, thus affecting the pattern of cell growth. Cells that never activated the NE (such as *rga4*Δ monopolar mother cells) produce one daughter cell that is born without any mark, causing symmetrical Cdc42 activation, and bipolar cell growth. To test this hypothesis genetically, we considered the possibility that a growth-dependent marker directs the organization of the Scd1/Scd2 GEF complex that activates Cdc42 at the cell tips. Because the organization and amplification of Scd1 at the cortex depends on the scaffold protein Scd2, we asked whether Scd2 remains at the tips during mitosis. We measured the intensity of Scd2-GFP at the cell tips of septated cells and found that while most Scd2-GFP is redistributed to the site of cell division, small amounts of Scd2-GFP can also be observed at the cell tips in WT and *rga4Δ* cells during cell division. This interesting finding suggests that Scd2 may indeed have a function in WT cells to promote Cdc42 amplification at the OE following mitosis. However, we found Scd2-GFP is similarly present at both tips in either *rga4*Δ monopolar or *rga4*Δ bipolar cells undergoing cell division, suggesting that the divergent Cdc42 dynamics observed in daughters of monopolar *rga4Δ* mother cells do not depend on asymmetric inheritance of the Scd2 complex.

The second hypothesis invokes the effects of asymmetric cell division of the monopolar mother cell, leading to daughter cells of different cell volumes. This hypothesis requires the bipolar daughter cell to be significantly larger than the monopolar cell. The cell born with a larger cell volume or larger surface area would then display initial conditions that favor bipolar growth, because it contains larger amounts of Cdc42 activators or a higher concentration of polarity factors at the cell cortex. Thus, we analyzed the dimensions of the two *rga4*Δ daughter cells. We found that the *rga4*Δ bipolar daughter cell is indeed larger than the monopolar *rga4*Δ daughter cell. Consistent with volume affecting the levels of Cdc42 activation, decreasing the volume of the bipolar cell by introduction of the *pom1*Δ mutation restores the normal, monopolar pattern of growth in young daughter cells that would otherwise be bipolar. However, the difference in volumes between *rga4*Δ monopolar and *rga4*Δ bipolar daughter cells at birth is rather small (approximately 20%), much smaller than the volume differences that our mathematical model requires to induce a significant change in the behavior of the system. Furthermore, we found that a small volume asymmetry is also sometimes observed in daughters of bipolar mothers that continue to grow in a monopolar way. Thus, these observations suggest that volume asymmetries are not sufficient to generate divergent Cdc42 dynamics in the *rga4*Δ daughter cells.

Finally, the third hypothesis implicates an asymmetric inheritance of Cdc42 regulators in *rga4*Δ daughter cells to explain divergent Cdc42 dynamics. Gef1 and Rga4 cooperate to regulate the active Cdc42 distribution at the cell ends (Das et al., 2015). Manipulating the levels of Gef1 and Rga4 controls cell diameter and bipolar cell growth (Das et al., 2015). This finding suggested that divergent growth patterns present in *rga4*Δ mutant cells may be a consequence of cells having different levels of Cdc42-GTP. Indeed, we found that loss of Cdc42 GEF *gef1* partially suppresses the divergent growth patterns observed in *rga4*Δ mutant cells, suggesting bipolar *rga4*Δ cells may have different initial levels of Cdc42 activity, as suggested by the model predictions of hypothesis three. However, we did not find evidence for asymmetrical localization of Cdc42 GEF Scd1 (which displays a similar localization as Scd2) or Cdc42 GEF Gef1 (which is normally not visible at cell tips (Das et al., 2015)) in *rga4*Δ newly born daughter cells, indicating that asymmetric inheritance of Cdc42 GEFs is likely not the reason for divergent Cdc42 dynamics.

In our SGA screen, we found that cells that lack *rga4* or both *rga4* and *rga6*, the daughter cell that lacks a previously growing end is born with a greater volume than the daughter cell that inherits a previously growing end. This asymmetry in cell volume is not due to misplacement of the septum along the center axis of the cell (not shown). Rather, monopolar *rga4*Δ cells and *rga4*Δ *rga6*Δ cells have a broader tip at the non-growing end than at the growing end, an effect that is more dramatic in the double mutant. This difference in tip shape alters the overall shape and surface area of the cell, which becomes broader near the nongrowing cell end. Thus, *rga4*Δ cells display an irregular cell shape, dividing asymmetrically even when the septum is placed in the middle of the center axis of the cell. This morphological asymmetry is amplified by loss of a second Cdc42 GAP protein, Rga6, suggesting that altered localization or inheritance of other Cdc42 GAP proteins may be altered in *rga4*Δ cells.

### A role for Cdc42 GAP proteins in linking the pattern of cell growth in the mother cell to Cdc42 dynamics in the daughter cells

Rga4 is a negative regulator of GTP-Cdc42, and its lateral localization helps to define a dynamic border that limits the extent of Cdc42 activation at the cell tips (Das et al., 2015). Consistent with this function, *rga4Δ* cells have broader distribution of active Cdc42 and a wider cell diameter (Kelly and Nurse, 2011). Rga6, a second Cdc42 GAP in fission yeast, cooperates with Rga4 in the control of Cdc42 dynamics, cell diameter, and polarized cell growth (Revilla-Guarinos et al., 2016). In the absence of *rga4*, we found that Rga6 localization becomes more asymmetrically localized towards the single growing cell tip in monopolar mother cells. This asymmetry reflects the pattern and history of cell growth and is maintained throughout mitosis. Thus, immediately following cell division, newly born *rga4*Δ cells display different levels of Rga6 at the cell membrane in a manner that depends on the pattern of growth of the mother cell. Loss of *rga4*, by altering the distribution of Rga6, further reduces the negative regulation of Cdc42 in cells that do not inherit a previously growing OE; in these cells, Cdc42 oscillations are immediately symmetrical, and cell growth bipolar (Fig. 9).

**Figure 9.**
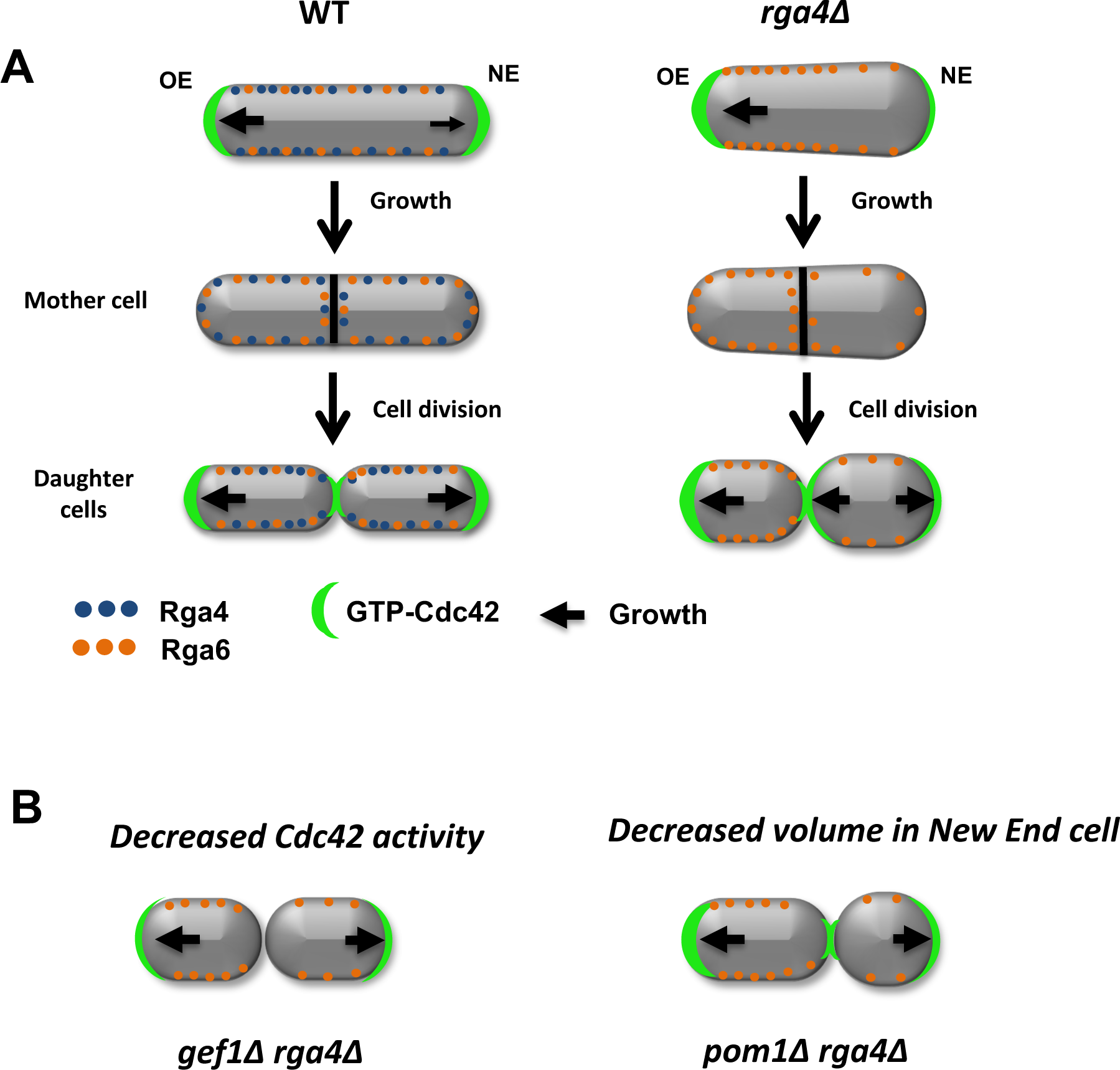
Role of Cdc42 GAP Rga4 in the initial state of Cdc42 activation in daughter cells. **A.** Cdc42 GAP Rga4 (blue dots) display a distribution with a bias towards the old growing end in wild-type cells (Das et al., 2007; Tatebe et al, 2008). In mutants that lack *rga4*, Cdc42 GAP Rga6 distribution becomes more asymmetrical in monopolar cells, resulting in daughter cells with different initial volumes and Cdc42 activity. Different levels of active Cdc42 between the two daughter cells, lead to divergent growth patterns. **B.** Mutations that decrease levels of active Cdc42 such as *gef1*Δ or decrease the volume of the larger daughter cell, such as *pom1*Δ, alleviate the growth pattern divergence of *rga4*Δ daughter cells.

Overall, our experimental observations support hypothesis 3 of our mathematical model, predicting how the presence of different amounts of Cdc42 regulators can generate different Cdc42 dynamics in *rga4*Δ daughter cells. Indeed, we have experimentally discovered an asymmetrical inheritance of Cdc42 GAP Rga6 in *rga4*Δ cells. Additionally, *rga4*Δ daughter cells display subtly different cell volumes, likely also an effect of altered Rga6 distribution, which may also contribute to divergent Cdc42 dynamics, as predicted by hypothesis 2 of our mathematical model. Thus, it is also possible that the *rga4*Δ phenotype derives by a combinatorial effect of asymmetric inheritance of Cdc42 regulators (hypothesis 3) and a subtle difference in initial cell volumes at the start (hypothesis 2).

Conversely, we did not find experimental evidence to support the idea of a tip-aging “marker” that promotes Cdc42 activation in daughter cells (hypothesis 1). Interestingly, we find mathematically that the presence of a tip-aging marker renders the system less able to switch to more symmetrical oscillations, when the time-dependent growth of a wild-type cell is modeled, in the presence of noise (Supplemental Fig. 5). This suggest that the presence of a “mark” could actually be detrimental to Cdc42 redistribution in the cell and impede bipolar growth activation. Indeed, the robustness of the third mathematical hypothesis is also supported by the fact that addition of noise reproduces the transient transitions to bipolar oscillations, sometimes observed in time lapse images of monopolar *rga4*Δ cells (Supplemental Fig. 5). Such transient transitions that do not result in permanent growth pattern change, together with all demonstrated differences in concentrations and size between *rga4*Δ daughter cells, indicate that the growth pattern of *rga4*Δ cannot be fully explained by *rga4*Δ daughter cells finding themselves into either monopolar or bipolar valleys of attraction of a coexistence region that maintains itself through the cell cycle (Cerone et al., 2012). We instead suggest that the co-existence region may be encountered at different growth stages for different daughter cells.

In conclusion, our analysis of *rga4*Δ cells, by genetic and mathematical approaches, uncovered a possible biological significance of bipolar growth activation (NETO) (Fig. 9, hypothetical model). Although NETO is not required for cell growth, we find that NETO may function to ensure that all cells are born with similar initial conditions, to enable consistent growth patterns among a population of cells. The activation of NETO and bipolar growth in the mother cell is instrumental to allow localization of Cdc42 GAPs around both cell ends to ensure symmetrical inheritance of these Cdc42 negative regulators in the daughter cells. Therefore, our work revealed an unexpected and important role for Cdc42 GAPs in maintaining consistent Cdc42 dynamics from generation to generation. Because the mechanisms of Cdc42 control are highly conserved in higher organisms, this work may contribute to our understanding of asymmetric cell division in higher eukaryotes, with implications for development, stem cell maintenance, and human disease.

## Materials and methods

### Strains and Cell Culture

All *S. pombe* strains used in this study are isogenic to the original strain 972 and are listed in Supplemental Table 1. Cells were cultured in yeast extract (YE) or in Edinburgh minimal medium (EMM) plus required supplements at 32°C or at 30°C and grown exponentially for at least eight generations before analysis. Standard techniques were used for genetic manipulation and analysis (Moreno et al., 1991).

### Fluorescence Microscopy and Image Analysis

Live cell imaging was performed using a Zeiss Axiophot microscope, (connected to an Orca-ER Hamamatsu cooled high-resolution digital camera, a shutter, and a MAC 5000 shutter controller box), or an Olympus fluorescence BX61 microscope (Center Valley, PA) (equipped with Nomarski differential interference contrast (DIC) optics, a 100X objective, a Roper Cool-SNAP HQ camera (Tucson, AZ), Sutter Lambda 10 + 2 automated excitation and emission filter wheels (Novato, CA), and a 175 W Xenon remote source lamp with liquid light guide), or an Olympus BX71 inverted microscope (equipped with DIC optics, a 150X objective, a DG4 rapid wavelength switcher and a Hamamatsu EM CCD camera). Images were acquired using either Intelligent Imaging Innovations SlideBook image analysis software (Denver, CO) or Micromanager (version 1.4.15) from the Ron Vale laboratory at the University of California in San Francisco. ImageJ1.48g software (National Institutes of Health) was used to measure fluorescence intensities.

Analyses of cell growth, CRIB-GFP intensities dynamics were performed by spreading 5 μL of exponentially growing cells on to 35-mm glass bottom culture dishes underneath a pre-made slab of YE or EMM with 0.6% agarose and 1 mM ascorbic acid. The cells were then incubated at room temperature (25°C) for 30 minutes to allow cells to adapt. Cells were then visualized from the culture dishes using a Zeiss Axiophot microscope or BX71 inverted microscope. Correlation of growth and CRIB-GFP signal was performed as described in Das et al, 2012. Cell growth was measured as the increase in cell length from the birth scar to either cell tip. The localization of Scd2-GFP and Pom1-GFP was analyzed using either a Zeiss Axiophot microscope or an Olympus fluorescence BX61 microscope. One milliliter of exponentially growing cells was pelleted, and the majority of the supernatant was removed leaving approximately 10µl. Then, 1 µl of cells was spread on to a 25 × 75mm 1.0mm thick microslide and covered with a 22 × 22mm micro cover glass. For Calcofluor staining, 1-2µl of Calcofluor solution (1mg/ml) was added to freshly pelleted cells, and the staining was visualized using epifluorescence.

### Mathematical Model

To model the polarization pattern of *rga4*Δ cells, we started from the model of (Das et al., 2012). We first present a brief summary of this model. For more details and for method of numerical solution see (Das et al., 2012). The model considers a population of a limiting component (that could represent Cdc42, Cdc42-GTP, or Cdc42 GEFs) distributed among three subpopulations: a population in the cell middle (cytoplasmic or membrane-bound with total number *C*_cyto_) and one at each tip (total numbers *C*_tip1_, *C*_tip2_). The total amount is *C*_tot_ = *C*_tip1_ + *C*_tip2_ + *C*_cyto_ and is assumed to increase in proportion to cell volume with constant rate *dC*_*tot*_/*dt*. The cytoplasmic population is considered to be well mixed because cytoplasmic or membrane-bound proteins typically diffuse across the cell over a shorter time compared to the time required for significant change of tip concentration (∼ minutes). The dynamical equations in (Das et al., 2012) were the same for each tip:

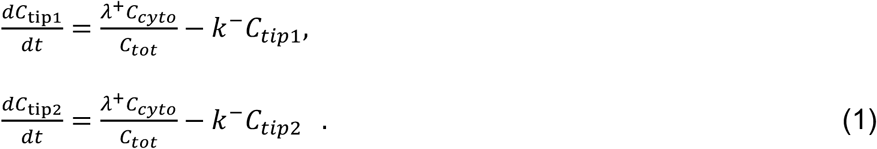

The factor *C*_*tot*_, which is proportional to cell volume, appears in the denominator of the association term because the cytoplasmic concentration interacts with the tip surface, a fixed area. Symmetry breaking was introduced through autocatalytic activation:

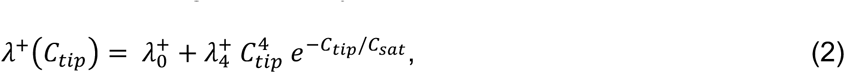

where 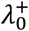 and 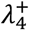 are constants. This nonlinear term represents positive feedback and generates asymmetry in short cells by allowing one tip to deplete the cytoplasmic pool and preventing the other tip from accumulating Cdc42. Saturation at level *C*_*sat*_ allows long cells to become bipolar. Asymmetric and symmetric oscillations result by assuming Cdc42 accumulation triggers its own removal (delayed negative feedback):

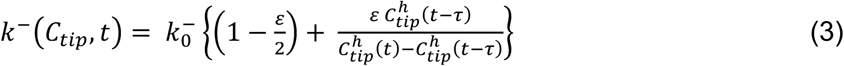

Here, 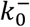 is a constant, *ε* determines the delayed dissociation strength, τ is delay time, and exponent *h* gives the nonlinearity of the effect.

As a simple mechanism to implement concentration fluctuations, Gaussian white noise of amplitude *γ* can be introduced to both *λ*^+^/*V* and *k*^−^ without allowing them to fall below zero. This method keeps mass conservation, and concentration amounts cannot become negative.

#### Model for Hypothesis 1

Cdc42 activation rate depends on the tip’s prior growth history. This model was implemented by introducing tip-aging dimensionless parameters *a*_1_ and *a*_2_, which describe how the rate of Cdc42 recruitment at either tip increases as a result of a history of prior growth. The following equations replace Equations (1):

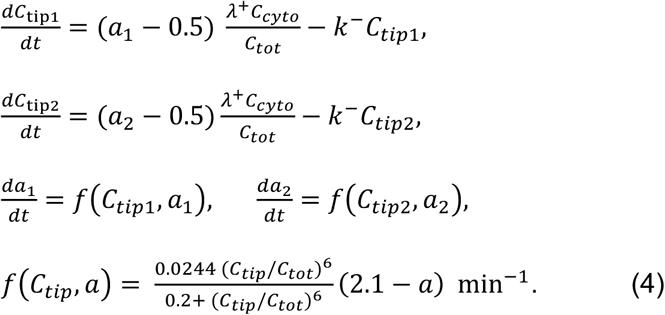

Here, both tips obey the same equations, but the aging parameter increases faster for the tip that happens to have the highest *C*_*tip*_, reaching up to a maximum plateau value assumed to be *a* = 2.1. A new tip formed at the division site is assumed to have a value *a*_*new*_ (= 1 for simulations of *rga4*Δ cells). The functional dependence of the rate of tip aging parameter increase *f* (*C*_*tip*_, *a*) is chosen such that the aging parameter can reach the plateau smoothly through the cell doubling time under conditions when the tip accumulates a significant fraction of Cdc42 over the cell doubling time, assumed to be 240 min. Each daughter is assumed to start with half of the volume of the mother. A daughter inherits one aged tip carrying the aging parameter of one the two mother tips and a new tip that starts with aging parameter *a*_*new*_.

#### Model for Hypothesis 2

Daughters of monopolar *rga4*Δ mothers inherit unequal volumes. The equations for this model are the same as the ones in (Das et al., 2012). We study the behavior of cells that start with different total amounts *C*_*tot*_, in proportion to the differences in volume.

#### Model for Hypothesis 3

Daughters of monopolar *rga4*Δ mothers inherit unequal Cdc42 regulators. This model is described by Equations (1)-(3) but we further assume that 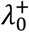 is a dynamic global parameter (i.e., has the same value for both tips and varies over time). This parameter relaxes towards a reference value 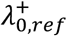 over the course of the cell doubling time:

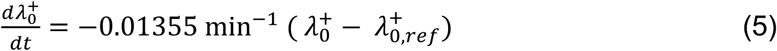

This model was explored as a function of the initial value of 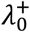 that is assumed to be different in the two daughters of monopolar *rga4*Δ mothers, each inheriting a 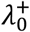 at birth above and below 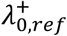, respectively.

### Model parameter values

In all cases, amount at tips and cell middle measured with respect to the saturation parameter in the model of wild type cells without tip aging, 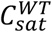. Model parameters for wild type cells are the same as (Das et al., 2012).

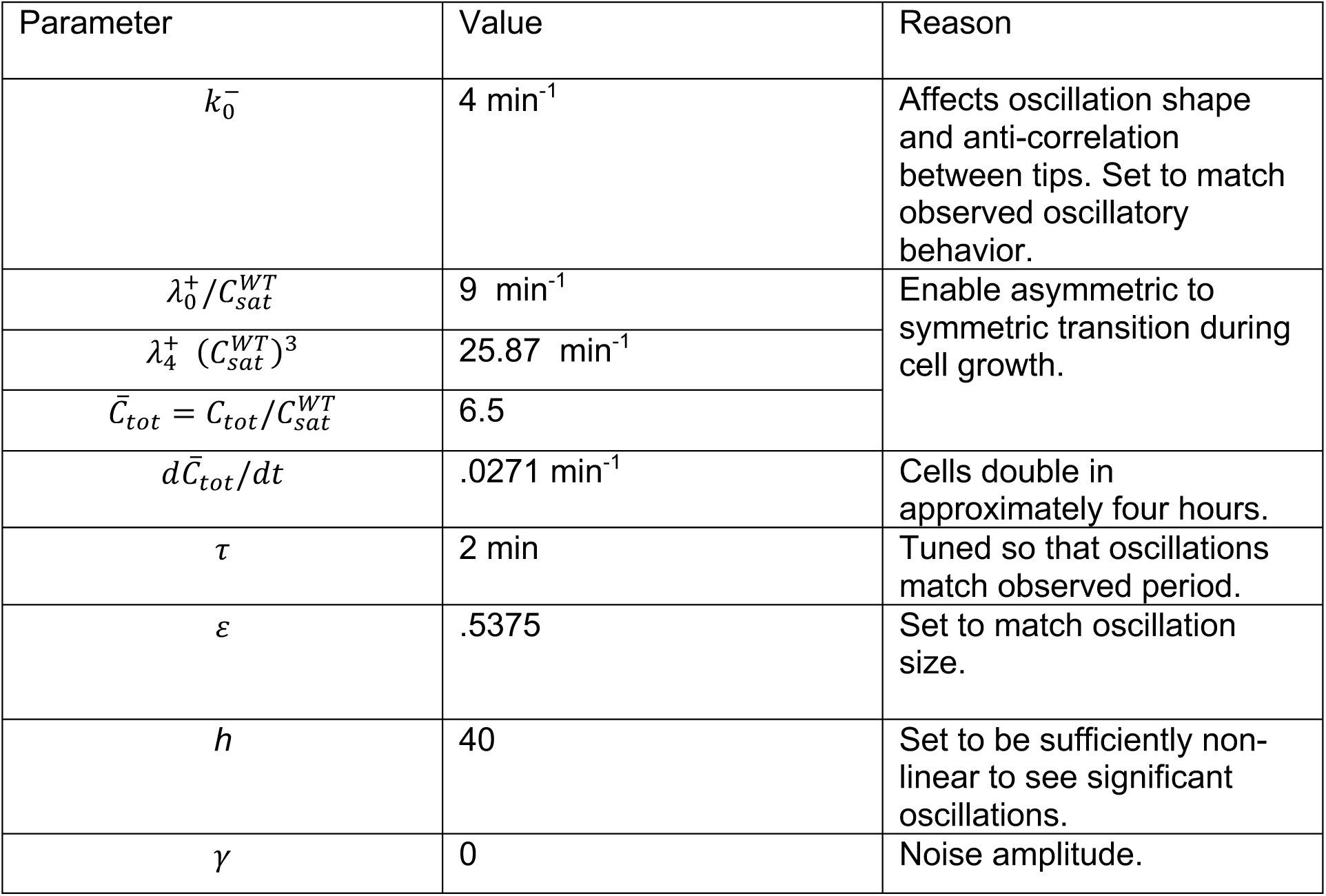

| Parameter | Value | Reason |
| --- | --- | --- |
| $k_0^-$ | 4 min<sup>-1</sup> | Affects oscillation shape and anti-correlation between tips. Set to match observed oscillatory behavior. |
| $\lambda_0^+ / C_{sat}^{WT}$ | 9 min<sup>-1</sup> | Enable asymmetric to symmetric transition during cell growth. |
| $\lambda_4^+ (C_{sat}^{WT})^3$ | 25.87 min<sup>-1</sup> | |
| $\bar{C}_{tot} = C_{tot} / C_{sat}^{WT}$ | 6.5 | |
| $d\bar{C}_{tot}/dt$ | .0271 min<sup>-1</sup> | Cells double in approximately four hours. |
| $\tau$ | 2 min | Tuned so that oscillations match observed period. |
| $\varepsilon$ | .5375 | Set to match oscillation size. |
| $h$ | 40 | Set to be sufficiently non-linear to see significant oscillations. |
| $\gamma$ | 0 | Noise amplitude. |

Modified parameters for *rga4*Δ cells, Hypothesis 1 (Fig. 3B; Supplemental Fig. 2A).

| Parameter | Value | Reason |
| --- | --- | --- |
| $C_{sat} / C_{sat}^{WT}$ | 1.6 | Effect of Rga4 deletion (see main text). |
| $\varepsilon$ | 1.85 | |
| $a_{new}$ | 1 | Low aging for new tips. |

Modified parameters for which model with tip aging history (Hypothesis 1) reproduces wild type polarity pattern (Fig. 3A; Supplemental Fig. 2A).

| Parameter | Value | Reason |
| --- | --- | --- |
| $C_{sat} / C_{sat}^{WT}$ | 1 | Close to value of model without tip aging. |
| $\varepsilon$ | 0.64 | |
| $a_{new}$ | 1.7 | Low aging for new tips. |

Modified parameters for *rga4*Δ cells, Hypothesis 2 (Fig. 3C).

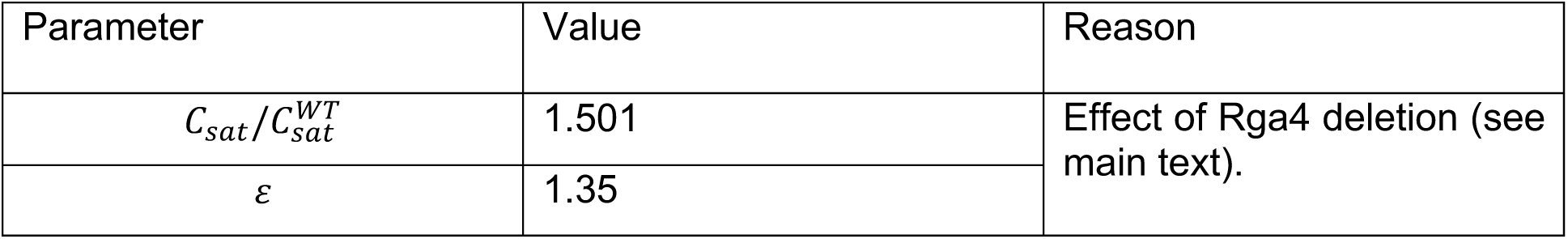

Modified parameters for *rga4*Δ cells, Hypothesis 3 (Fig. 3D; Supplemental Fig. 2B).

Modified parameters for *rga4* $\Delta$ cells, Hypothesis 3 (Fig. 3D; Supplemental Fig. 2B).
| Parameter | Value | Reason |
| --- | --- | --- |
| $C_{sat}/C_{sat}^{WT}$ | 1.6 | Effect of Rga4 deletion (see main text). |
| $\lambda_{0,ref}^+/C_{sat}^{WT}$ | 20 min<sup>-1</sup> | |

## Acknowledgments

This work in Dr. Fulvia Verde’s laboratory is supported by the National Institutes of Health R01 grant number GM129514. D.V. was supported by National Institutes of Health R01 grant number grant number R01GM114201. We thank Dr. Robert Tams for critical comments on the manuscript.

## Author Contributions

Conceptualization, M.R-P., I.N., M.D., D.V. and F.V.; Methodology, M.R-P., I.N., C.C., D.V. and F.V.; Investigation, M.R-P., I.N., C.C., D.W., G.D., P.B.; Formal Analysis, M.R-P., I.N., C.C., D.W., P.B., D.V., and G.D.; Writing – Original Draft, M.R-P. and I.N.; Mathematical model section: D.V.; Writing – Review & Editing: F.V.; Funding Acquisition: F.V.; Supervision: F.V.

