## Supplemental Figures for "Cdc42 GTPase Activating Proteins (GAPs) Maintain Generational Inheritance of Cell Polarity and Cell Shape in Fission Yeast"

A

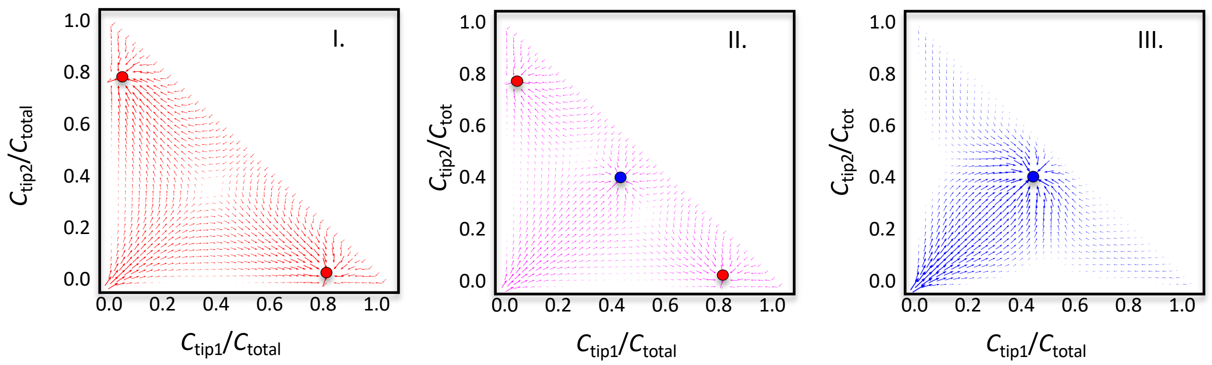


B

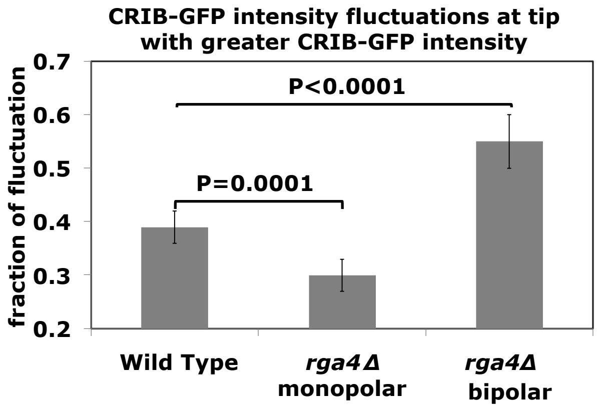


**Supplemental Figure 1. Mathematical model describing symmetric and asymmetric Cdc42 distribution depending on cell size.** A. Representative stream plots derived from the mathematical model describing Cdc42 oscillatory dynamics. The detailed model has been previously published in Das *et al.* 2012. $C_{tip}/C_{total}$ refers to the fraction of tip-bound Cdc42 ($C_{tip}$) relative to the total amount of Cdc42 in the cell ($C_{total}$). Arrows indicate the direction of progression of the system over time, and the length of the arrows corresponds with the speed along that path. Red circles indicate asymmetric stable steady states. Blue circles indicate symmetric stable steady states (attractors) of the system. For cells with small volume only asymmetric steady states are available (I). A coexistence region of asymmetric and symmetric steady states emerges as cell volume increases (II). In cells with a large volume only symmetric steady states are possible (III). B. CRIB-GFP dynamics at the cell tips in *rga4*Δ divergent daughter cells. Fluctuations of active Cdc42 are different in monopolar or bipolar *rga4*Δ cells. CRIB-GFP intensity was measured at tips in WT and *rga4*Δ cells, following cells in time by time-lapse microscopy. The relative fraction of active Cdc42 fluctuation at the cell tips was calculated as a ratio between the average intensity variation from maximal to minimal, and the average maximal peak of intensity, as observed during the time of the recording.

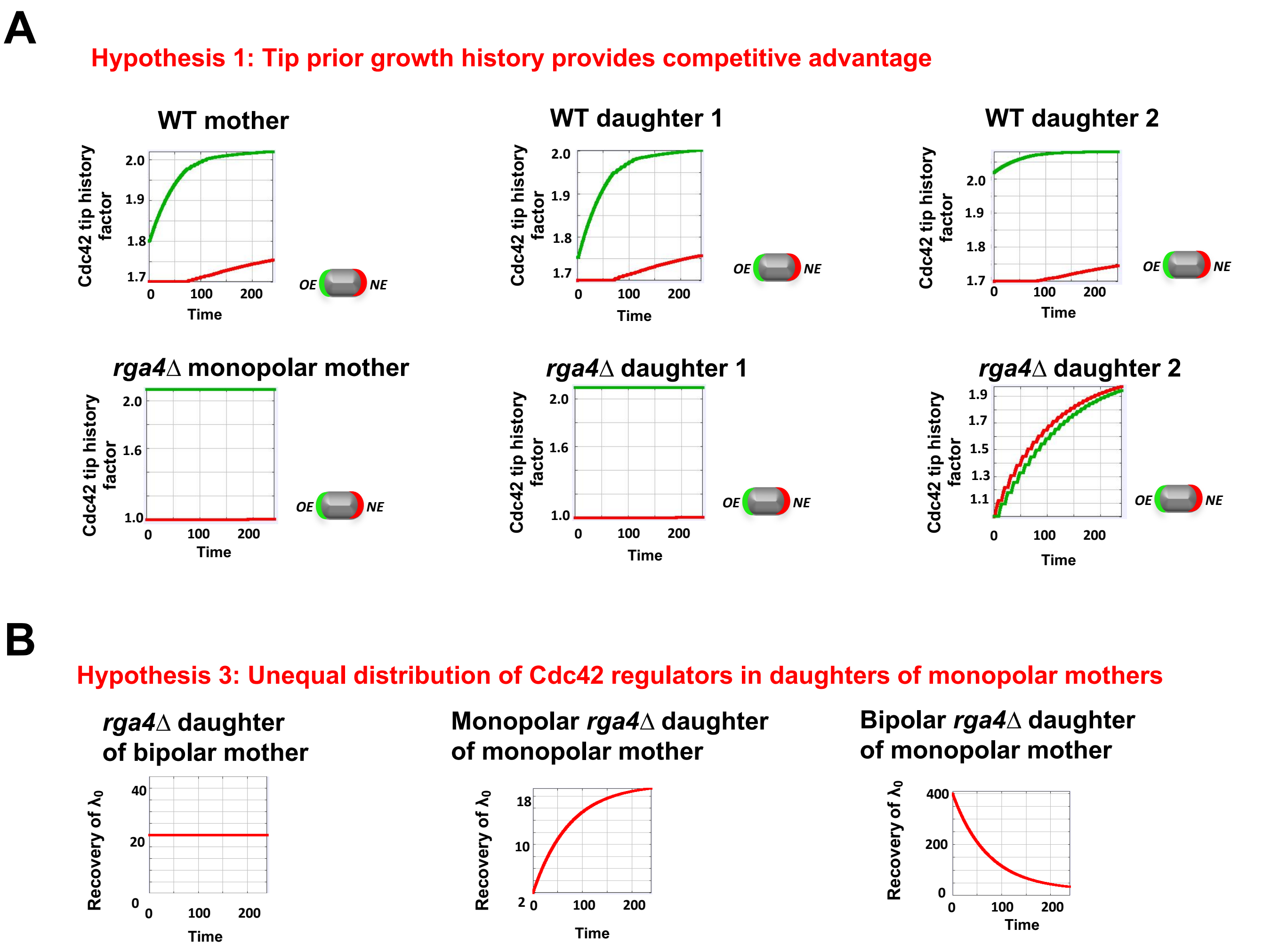


**Supplemental Figure 2. Mathematical models reproducing the *rga4*Δ phenotype. A.** In WT cells, the <u>aging parameter</u> (Hypothesis 1) can reach the plateau smoothly as the tip accumulates a significant fraction of Cdc42 over the cell doubling time, assumed to be 240 min. Each daughter is assumed to start with half of the volume of the mother. By contrast, in the case of *rga4Δ,* the aging is only restored for bipolar cells (daughter 2) **B.** Daughters of monopolar *rga4*Δ mothers inherit <u>unequal Cdc42 regulators</u> (Hypothesis 3). $\lambda_0^+$ is a dynamic global parameter that relaxes towards a reference value $\lambda_{0,ref}^+$ over the course of the cell doubling time. The initial value of $\lambda_0^+$ is assumed to be different in the two daughters of monopolar *rga4*Δ mothers, each inheriting a $\lambda_0^+$ at birth above and below $\lambda_{0,ref}^+$ , respectively.

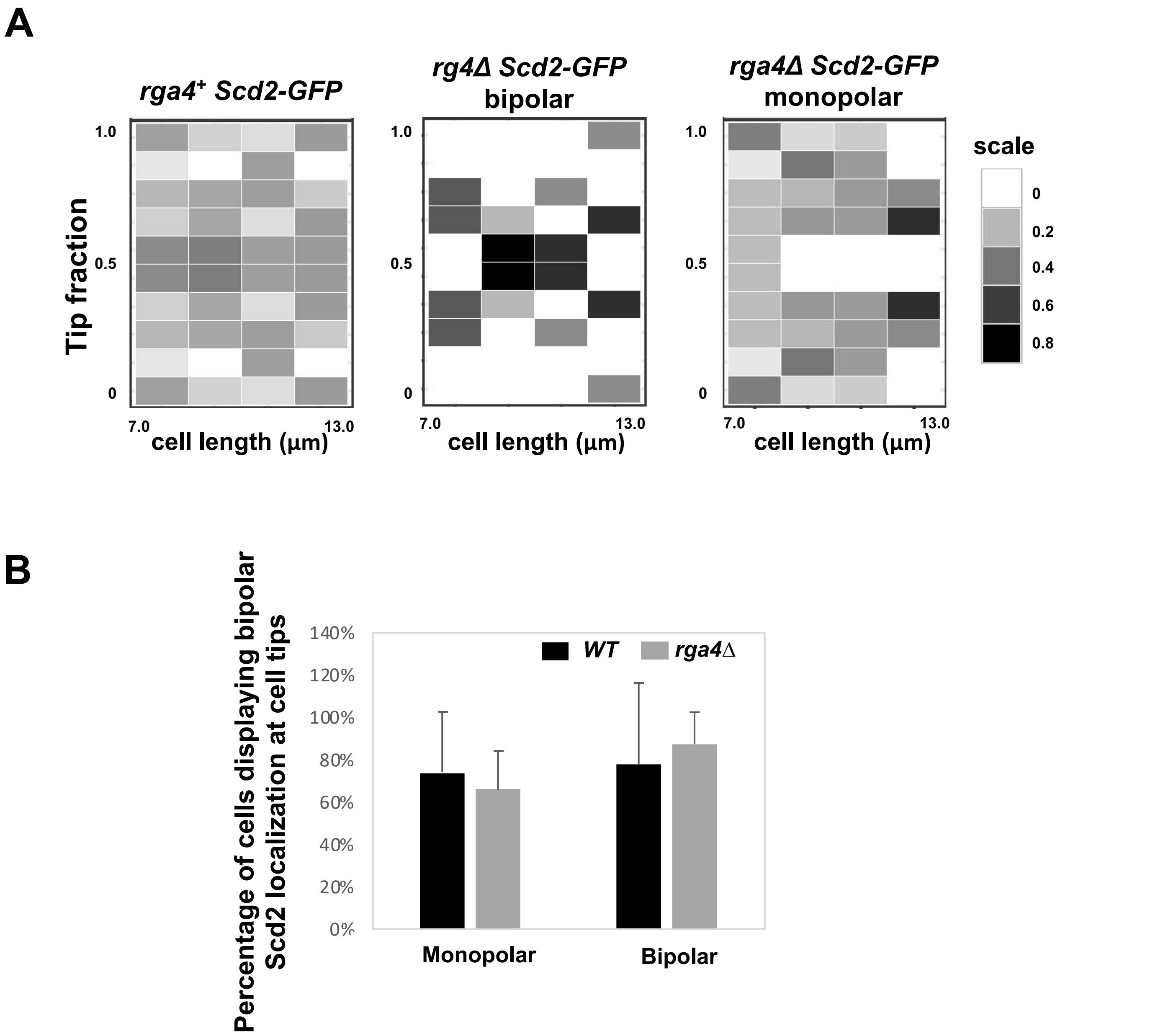


**Supplemental Figure 3**. **Scd2-GFP tip distribution in *rga4*Δ mutant cells. A.** Heatmap of Scd2-GFP tip fractions in growing, interphase wild-type (N=41) and *rga4*Δ bipolar (N=13) and monopolar cells (N=36). **B.** Scd2-localization at the cell tips during cell division in wild-type and mutant *rga4*Δ monopolar and bipolar cells (percentage of either total monopolar or total bipolar cells is shown, where Scd2 is visible).

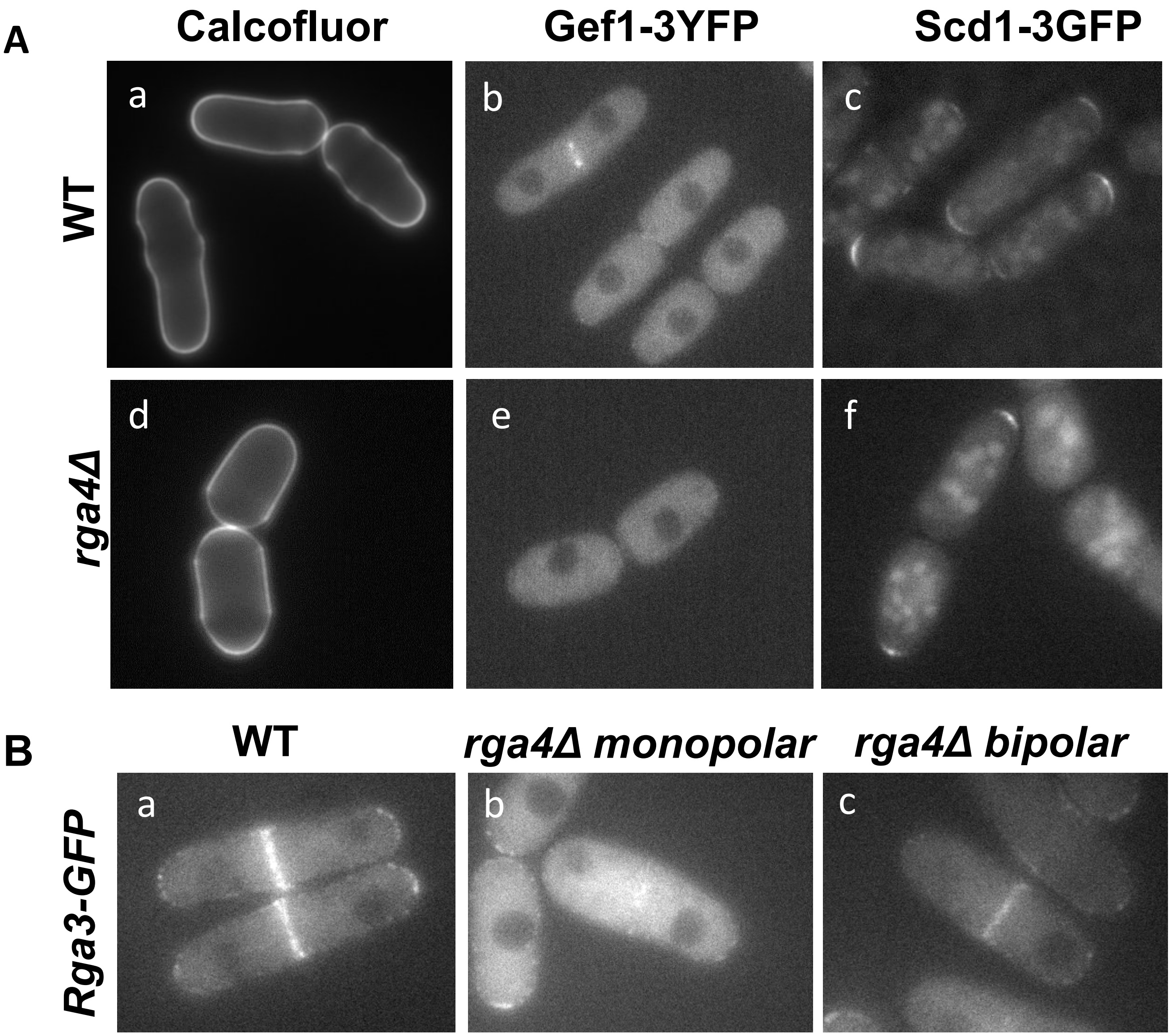


**Supplemental Figure 4. A. Cdc42 GEF Scd1 and Gef1 tip distribution in WT and *rga4*Δ mutant cells.** (a,d) Calcofluor staining in WT and *rga4*Δ cells. (b, e) Gef1 localization in dividing WT and *rga4*Δ cells. (c, f) Scd1 localization in dividing WT and *rga4*Δ cells. **B. Cdc42 GAP Rga3 tip distribution in WT and *rga4*Δ mutant cells.** a, WT. b, monopolar dividing *rga4*Δ cell. c, bipolar dividing *rga4*Δ cell.

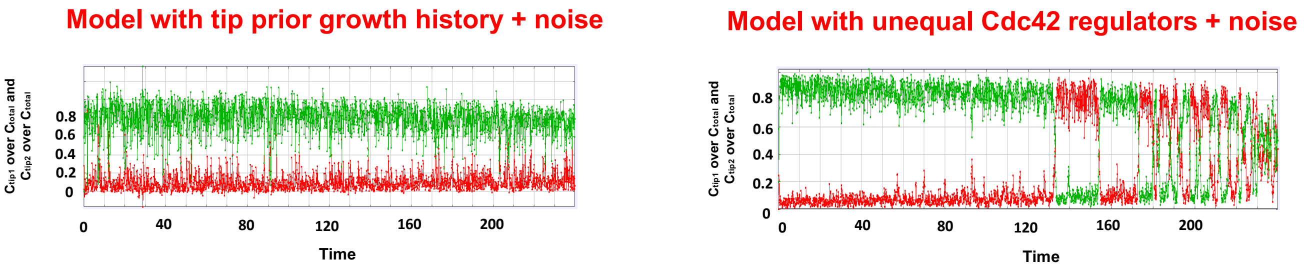


**Supplemental Figure 5**. Model behavior according to hypothesis 1 (tip growth history) or hypothesis 3 (unequal distribution of Cdc42 regulators), when noise is added as was performed in (Das et al., 2012).

**Supplemental Table 1.** List of strains used in this study.

| Strain | Genotype | Origin |
|---|---|---|
| *972* | *h⁻* | P. Nurse |
| *567* | *h⁻ ade6-704 leu1-32 ura4-D18* | P. Nurse |
| *CA5931* | *h⁻ shk1 promoter-ScGic2-CRIB-3xGFP:ura4⁺ ura4-294 ade-704 leu1-32* | Tatebe *et al.* 2008 |
| *FV1529* | *h⁻ ∆rga4::ura+ ade6-704 leu1-32 ura4-D18* | Das *et al.* 2007 |
| *FV1174* | *∆rga4::ura+ shk1 promoter-ScGic2-CRIB-3xGFP:ura4⁺ leu1-32 ura4-D18* | Das *et al. 2012* |
| *FV2401* | *Scd2-GFP-kanMX6 rlc1-tdTomato-natMX6 ade6-M21X leu1-32 ura4-D18 his3-D1* | This study |
| *FV2414* | *∆rga4::ura+ Scd2-GFP-kanMX6 rlc1-tdTomato-natMX6 ade6-MX21 leu1-32 ura4-D18 his3-D1* | This study |
| *SPAC24H6.09* | *∆gef1::kanMX4 ade6-M210 ura4-D18 leu1-32* | Kim *et al.*, 2010 |
| *FV1928* | *∆gef1::kanMX4 ∆rga4::ura+ ade6-M210 ura4-D18 leu1-32* | This study |
| *FV1381* | *h+ ∆gef1::ura+::gef1-3YFP-kanMX6 ade-704 leu1-32 ura4-D18* | Das, *et al.* 2015 |
| *FV1446* | *∆rga4::ura+ ∆gef1::ura+::gef1-3YFP-kanMX6 ade-704 leu1-32 ura4-D18* | This study |
| *SPBC354.13* | *∆rga6::kanMX4 ade6-M216 ura4-D18 leu1-32* | Kim *et al.*, 2010 |
| *FV2379* | *∆rga6::kan ∆rga4::ura+ ade6-M216 ura4-D18 leu1-32* | This study |
| *FV2461* | *∆rga6::nat rga6-3YFP-kanMX6 ∆rga4::ura+ ura4-D18 leu1-32* | This study |
| *FC1162* | *h⁻ pom1-GFP:kanMX* | Padte *et al.* 2006 |
| *FV2570* | *∆rga4::ura+ pom1-GFP:kanMX leu1-32 ura4-D18* | This study |
| *FV2567* | *rga4-mCherry-kanMX6 Cdr2-GFP-kanMX6* | This study |
